# panREPET: a reference-free pipeline for detecting shared Transposable Elements from pan-genomes to retrace their dynamics in a species

**DOI:** 10.1101/2024.06.17.598857

**Authors:** Somia Saidi, Mathieu Blaison, María del Pilar Rodríguez-Ordóñez, Johann Confais, Hadi Quesneville

## Abstract

The role of transposable elements (TEs) in host adaptation has gained lots of interest in the recent past. Individuals of the same species undergo independent TE insertions, causing genetic variability upon which natural selection can fosters adaptation of individuals to their environment. While *de novo* assembled genomes are becoming increasingly affordable, overcoming bias introduced by a single reference genome, suitable pangenomic approaches are required to explore genomes of a species. We developed a new pipeline called panREPET that identifies TE insertions shared by groups of individuals. Unlike other pangenomic tools, panREPET operates independently of a reference genome and provides the precise sequence and genomic coordinates of each TE copy for each genome. We showcase here the potential of this tool on TE insertions shared among 42 *Brachypodium distachyo*n genomes and compared our results against existing tools to demonstrate its better efficiency. With this tool, we were able to date two major TE bursts corresponding to major climate events: 22 kya during the Last Glacial Maximum and 10 kya during the Holocene, showing a potential link between environmental stress and TE activity.

## Introduction

Transposable elements (TEs) are mobile DNA elements that can invade genomes by transposition. Their insertions may be neutral to their host or deleterious, for instance, by disrupting gene expression (Lish *et al*., 2013 ; Hirsh *et al*., 2017 ; Bourque *et al*., 2018). Host defense may involve epigenetic regulation (Lish *et al*., 2013) or purifying selection to purge deleterious TE (Langmüller *et al*., 2023). TE insertions may also be beneficial in fostering genome evolution through introduction of functional novelties (Schrader *et al*., 2019). In natural populations, TE insertion polymorphism frequency is subject to drift, selection and population migrations (Bourgeois *et al*., 2019). Understanding how TEs can enable a species to adapt to a local environment requires a detailed study of their insertions in genomes of different individuals. At the species level, the pangenomic approach seeks to detect intra-specific diversity of TEs and to retrace their dynamics. A TE pangenome can be described by (i) TE insertions present in all individuals of the species (core-genome), (ii) insertions present only among a subset of individuals and (iii) individual-specific insertions. Some TEs are known to be sensitive to abiotic stress and can, therefore, affect nearby genes in response to environmental conditions (Le TN *et al*., 2014 ; Makarevitch *et al*., 2015; Schrader *et al*., 2019. Using a pangenomic approach, studies have shown that recent TE insertions, at the intra-specific level, are involved in local adaptation. For example, a population genomic study of the *Arabidopsis thaliana* natural accessions revealed extensive intraspecific variation in *ATCOPIA78* copies, which correlates with temperature variation (Quadrana *et al*., 2016). A study in the rice suggested that TE activation might be induced by external stimuli instead of resulting from alterations of genetic factors involved in TE silencing pathways (Carpentier *et al*., 2019).

Detecting TE polymorphisms is challenging because of their repeated nature but also for detecting TE insertions absent from the reference when TE pangenome studies are based on sequence reads alignments from different genomes of the species to an assembled reference genome. Moreover, regions absent from the reference genome are unavailable for analysis as not assembled. Nowdays, several *de novo* assembled genomes for the same species are becoming available. Then, we developed a new pipeline, called panREPET, that do not depend on a reference genome to study TE insertions in a species. panREPET overcomes the reference dependency by comparing TE copies between each pair of *de novo* assembled genomes. panREPET can also facilitate dating of TE dynamics at the species level for all types of TE sequences.

To showcase the potential of panREPET, we tested it on *Brachypodium distachyon* for which many genome sequences are available. Interestingly, this annual Mediterranean plant, which has emerged as a grass genomic model, is undomesticatedand would allow to investigate TE dynamics in the context of adaptation to a mosaic of Mediteranean climates (Hasterok *et al*., 2022). Here, we used 54 *de novo* whole-genome assemblies available *for B. distachyon* (42 after filtering for best assembly qualities) (Gordon *et al*., 2017). Those accessions, initially sequenced to perform the *B. distachyon* pangenome, occur in contrasting ecological niches, are genetically diverse and represents three (B_West, B_East and A_East) of the five genetic clades since then identified in this species (Minadakis *et al*., 2023). In contrast to more recently sequenced accessions from Italy and the Balkans, the high-coverage employed by Gordon *et al*. 2017 (median genome coverage of 92x, mean assembled genome size of 268 Mbp, which is very close to the 272 Mbp reference genome size) allowed *de novo* assemblies. Because each accession has been annotated for genes but also used to investigate TE dynamics with short-read and population genomic approaches (Minadakis *et al.,* 2023 ; Horvath *et al*., 2023), they constitute an ideal dataset to test and benchmark panREPET. The questions on *B. distachyon* TE that we seek to address here to illustrate panREPET potential are: what are the main transposition events in the species, which TE families transposed, and when did it occurFor the oldest insertions, the question is what are the mechanisms of their conservationFor bursts of recent transpositions in few individuals, the question is whether external factors have contributed to their activation ?

Here, we will first present the panREPET pipeline and will compare its results to alternative tools used in recent TE pangenomic studies to demonstrate its performances. In a second part, we will show an application illustration to 42 genomes from *B. distachyon*. We describe the TE evolutionary history by estimating transposition bursts events precisely. We found co-occurence of the date of these events with climate changes that occurred in the past. Therefore, we will show that climate factors may, in some cases, explain TE dynamics.

## Results

### panTEannot : a TEannot adaptation

The TE reference library was built from the Bd21 v3.2 genome sequence using the TEdenovo pipeline from the REPET package (Flutre *et al*., 2011) (see Materials and Methods - Genome analysis tools). Our pangenomic approach aims to capture as much diversity as possible from all genomes. Consequently, we annotated each genome for its TE content independently using TEannot with the Bd21 TE reference library (**Fig. 1a**). But, this TE annotation process is time-consuming and requires a fast method for serial annotation of the 54 genomes (**Table 1**). Hence, we modified the TEannot pipeline from the REPET package (Quesneville *et al*., 2015) into a lighter version. Specifically, as we are interested in inter-individual TE variability, which is due in particular to recent TE transpositions, we focused on non-degenerate sequences, i.e. complete TE copies as a proxy. In the TEannot pipeline of REPET, the similarity search step is done by Blaster (Quesneville *et al*., 2003), Censor (Jurka *et al*., 1996) and RepeatMasker (Smith *et al*., 2013-2015) to maximize the sensitivity. As complete TE copies are easy to detect, there is no need to maximize the sensitivity. Hence, we kept a single tool, Blaster, and removed Censor and RepeatMasker. With the same rationale, we also removed SSR search, Spurious hits removal and Fragment connection (TEannot steps 4, 5 and 7) as they have low impact on detection of complete TEs. To improve parallel computation, we implemented it with the Snakemake framework (Mölder *et al*., 2021) removing MySQL dependencies and the use of a computer cluster job scheduler such as Sun Grid Engine or Slurm. This new implementation is called panTEannot. As we aim to keep good sensitivity and specificity with faster method, we benchmarked its performance by comparing panTEannot to TEannot from REPET v3.0. We performed a complete TE annotation with TEannot from REPET v3.0 on the genome sequence reference Bd21 to be used as the golden standard. We also performed annotation with TEannot REPET v3.0 run without Censor and RepeatMasker, to mimic analysis by panTEannot. We compared the different TE annotations against the golden standard at the nucleotide level (**Table 1**). We calculated sensitivity and specificity as in Baud *et al*. 2019 (see Materials and Methods - Genome analysis tools). As expected, TEannot from REPET 3.0 has a better sensitivity (0.9985) than panTEannot (0.8951). However panREPET has a better specificity (0.9267 against 0.8979) and is faster. As in a pangenomic context, we are more interested in detecting true positives than all potential copies including very old and thus degenerate ones, we considered this result as a satisfactory compromise.

**Figure 1:**
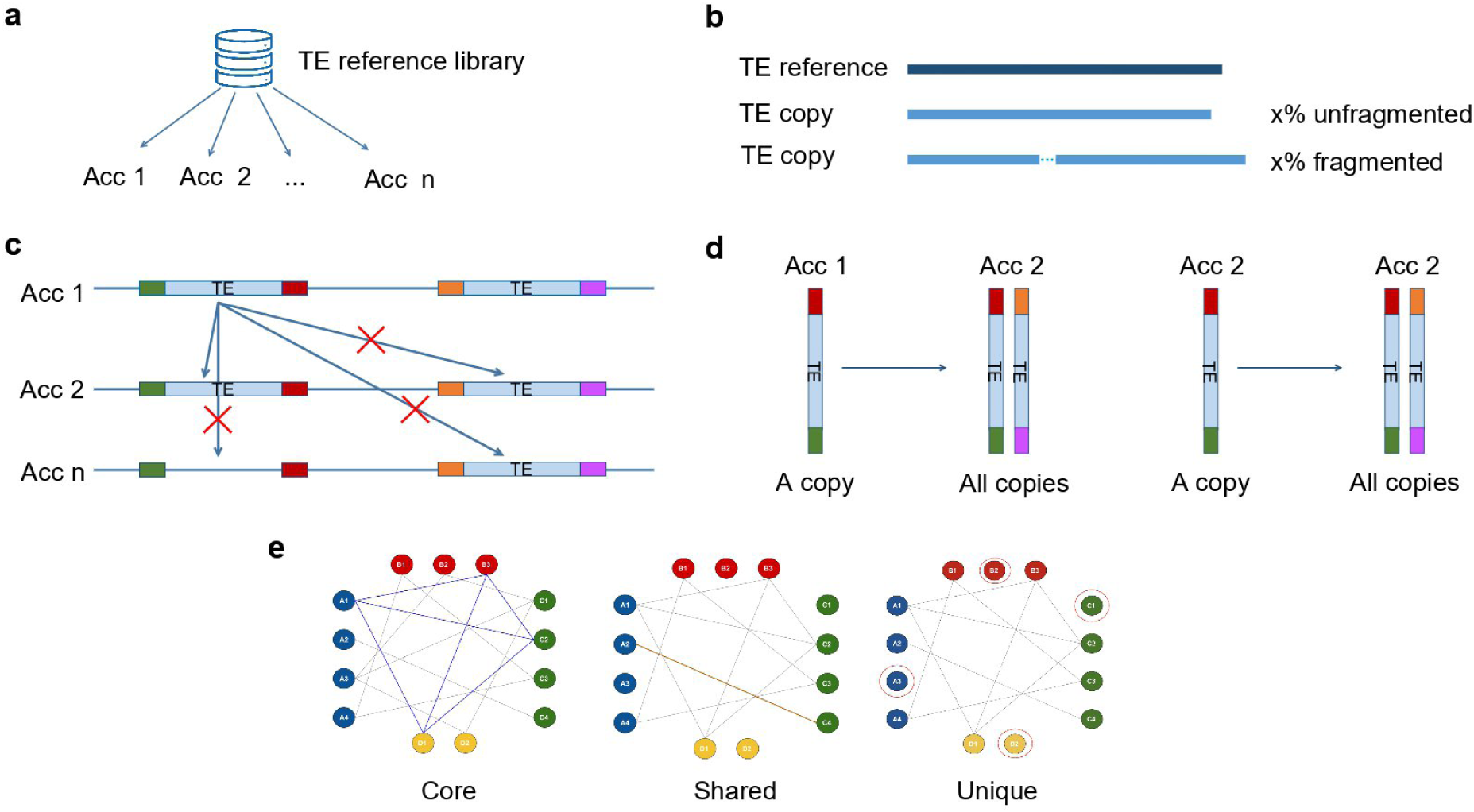
panREPET steps. **a** Independent TE genome annotation with panTEannot using a TE reference library. **b** Extraction of copies according to their coverage percentage in the alignment with their reference (e.g. *x* is between 95% and 105% for complete copies). **c** A TE copy from accession 1 is matched with TE copies having the same flanking regions in order to detect copies located at the same locus. **d** Bidirectional best-hit detection. **e** Clique extraction to identify TE insertion. Copy type identification: core, shared, or unique. A circle represents a TE copy, a color represents an accession, and a bold line represents a bidirectional best-hit detection. Acc: accession.

**Table 1:**
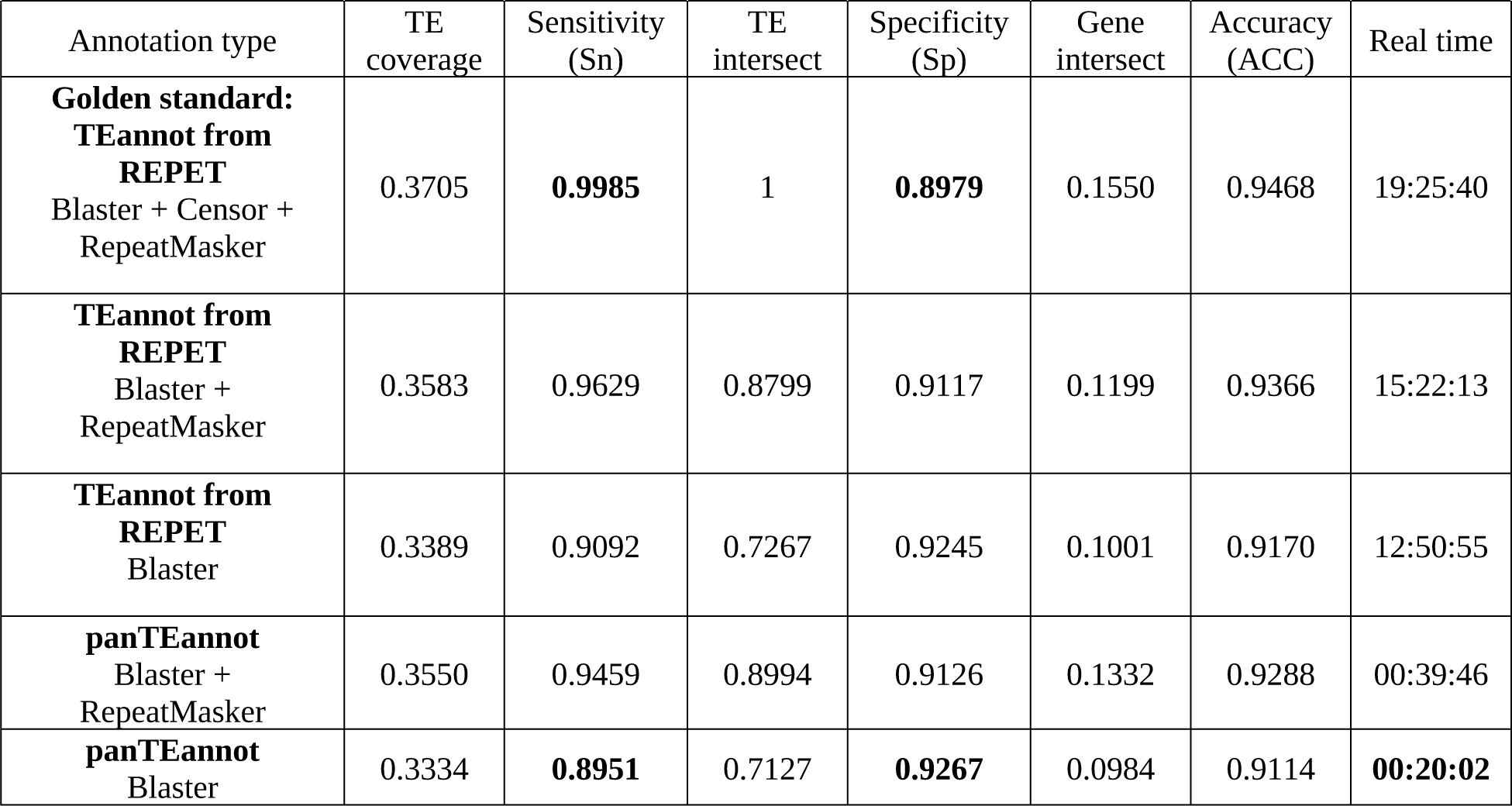
Comparison of performances of TEannot from REPET 3.0 with panTEannot on Bd21 genome sequence v3.2. Real times were obtained on a 32 core 64 RAM virtual machine.

| Annotation type | TE coverage | Sensitivity (Sn) | TE intersect | Specificity (Sp) | Gene intersect | Accuracy (ACC) | Real time |
| --- | --- | --- | --- | --- | --- | --- | --- |
| **Golden standard:**<br>**TEannot from REPET**<br>Blaster + Censor + RepeatMasker | 0.3705 | **0.9985** | 1 | **0.8979** | 0.1550 | 0.9468 | 19:25:40 |
| **TEannot from REPET**<br>Blaster + RepeatMasker | 0.3583 | 0.9629 | 0.8799 | 0.9117 | 0.1199 | 0.9366 | 15:22:13 |
| **TEannot from REPET**<br>Blaster | 0.3389 | 0.9092 | 0.7267 | 0.9245 | 0.1001 | 0.9170 | 12:50:55 |
| **panTEannot**<br>Blaster + RepeatMasker | 0.3550 | 0.9459 | 0.8994 | 0.9126 | 0.1332 | 0.9288 | 00:39:46 |
| **panTEannot**<br>Blaster | 0.3334 | **0.8951** | 0.7127 | **0.9267** | 0.0984 | 0.9114 | **00:20:02** |

On the 54 genomes, panTEannot took 48 hours on a 32 core 64 Gb RAM virtual machine which we consider to be a good performance. The average TE sequence coverage percentage of the genome sequence is 29.9%. Unassembled genome parts correspond mostly to TEs due to their repetitive content. We observed that the TE coverage depends on the assembly quality: genomes with a TE coverage percentage below 30% are characterized by an assembled genome size after removing ‘N’ occurrences lower than 95% of the reference genome (256 Mbp) **(Supplementary Table 1)**. This threshold led us to remove from our study ABR8, Bd3_1, Bd30-1, BdTR11a, BdTR12c, BdTR5i, BdTR8i, Gaz-8, Koz_3, Mur1, Tek-2 and Tek-4. Note that the BdTR11a genome is larger than 256 Mbp (about 266 Mbp) but presents only a TE sequence coverage percentage of 19%. Another useful metric for TE detection quality is the contig N50. The longer the contigs are, the better TEs are detected. Hence, we removed BdTR11a because in addition to not having a large assembled genome size, its contig N50 is low compared to others (17 kbp against 24 kbp in median).

### panREPET pipeline

We developed the panREPET pipeline to identify shared TE insertions between genomes by comparing them pairwise. We divided the pipeline into the following steps:

#### Step 1: Extract TE copies according to coverage percentage in the alignment between the sequence TE copy and its reference

The coverage percentage between the TE copy and its reference sequence from the TE library is a parameter (covcons parameter), allowing extraction of either complete or partial TE copies (**Fig. 1b**).

#### Step 2: Add flanking genomic sequence

In order to recognize copies that are located at the same genomic locus, we retrieve their flanking sequences by simple coordinate extension on both sides using *bedtools slop* from the Bedtools package (Bioinformatics, 2024) (**Fig. 1c**).

#### Step 3: Bidirectional best-hit detection

Bidirectional best-hit detection first consists of aligning TE copies extended by their flanking sequences between accessions to find their best hits and vice versa (**Fig. 1d**). Only couples of reciprocal best hits are retained. This approach was inspired by methods for orthologous gene detection (Emms *et al*., 2015). Pairwise comparisons are performed by Minimap2 (Li, 2018), followed by Matcher (Quesneville *et al*., 2003), which merges nearby colinear alignment fragments. Furthermore, panREPET offers the possibility of comparing only TE copies present on the same chromosomes between accessions, that permits to increase the computational efficiency and reduce false positives. It is then also possible to highlight shared TE insertions that have undergone an inter-chromosomal translocation event or cases of segmental duplication corresponding to paralog copies (see Discussion).

#### Step 4: From bidirectional best hits to TE insertions

Pairs of TE copies identified in step 3 are used to build an undirected graph where the TE copies are nodes and where edges identify copies found in reciprocal best hits. In graph theory, a clique is a subset of nodes in a graph in which every node is connected to every other node in the clique. We consider a clique as a proxy of a unique TE insertion found at the same locus in several genomes. Cliques are found using NetworkX (Hagberg *et al*., 2008) version 2.4. The method *read edgelist* from NetworkX constructs the graphs that associate pairs of copies through bidirectional best hits. Then, the method *find_cliques* from NetworkX detects all maximal cliques in the constructed graphs. For each node *n*, a maximal clique for *n* is a largest complete subgraph containing *n*. Note that a shared TE insertion can correspond to several cliques when at least two accessions in the subset of accessions do not success the bidirectional best hits detection step. panREPET gives in the final output the multiple possibilities.

Finally, each TE copy of each accession is classified on core, shared or unique if they are found respectively in all individuals, some of them, or in only one (**Fig. 1e**). Adding or deleting a genome will not require the pipeline to be restarted from the beginning, since pairwise genome comparisons are independent of clique building.

### Retracing shared TE insertions

We used panREPET to identify shared TE insertions within *B. distachyon* genomes and compare the results to the literature. **Fig. 2a, 2b** represent a histogram of TE insertion number according to the number of accessions sharing the insertion based on our reliable set of 42 accessions. According to its shape (**Fig. 2a, 2b)**, we considered five pangenomic compartments: 1. Unique-TE insertions; 2. Cloud-TE insertions shared by 2-7 accessions (2–20%); 3. Shell-TE insertions shared by 8-38 accessions (20-95%); 4. Softcore-TE insertions shared by 39-41 accessions (95-98% of accessions) and 5. Core-TE insertions shared by all accessions. A unique-TE insertion suggests a new TE insertion in a single individual or a horizontal transfer. A cloud or shell or soft-coreTE insertion may correspond to an insert inherited from a common ancestor in some related individuals. Core-TE insertions are fixed inserts at the species level. panREPET identified 319,969 TE copies that cover their consensus over more than 95% (covcons = 95-105%). Among them, it detected 18,513 shared TE insertions including 595 core (1.2%); 1,139 soft-core (2.4%); 9,125 shell (19.4%) and 7,654 cloud (16.2%). 28,497 are unique TE insertions (60.6%). The vast majority of TE insertions appear to be unique (**Fig. 2a, 2b)**. The pipeline ran for 20 hours on a 64 core 128 Gb RAM virtual machine. By including partial copies (covcons = 75-125%), it extracted two times more copies (653,695) corresponding to 35,922 shared TE insertions and 82,347 unique insertions. We thus observed a higher proportion of core (1.4%) and unique TE insertions (69.6%) (**Fig. 2b**). Longer TE copies tend to be unique or present in a small number of individuals (**Fig. 2c**). The pipeline ran for 3.6 days on a 64 core 128 Gb RAM virtual machine. Not extracting all TE copies by setting the coverage between copies and their consensus parameter (covcons) aims to save computational time, especially for huge genomes.

**Figure 2:**
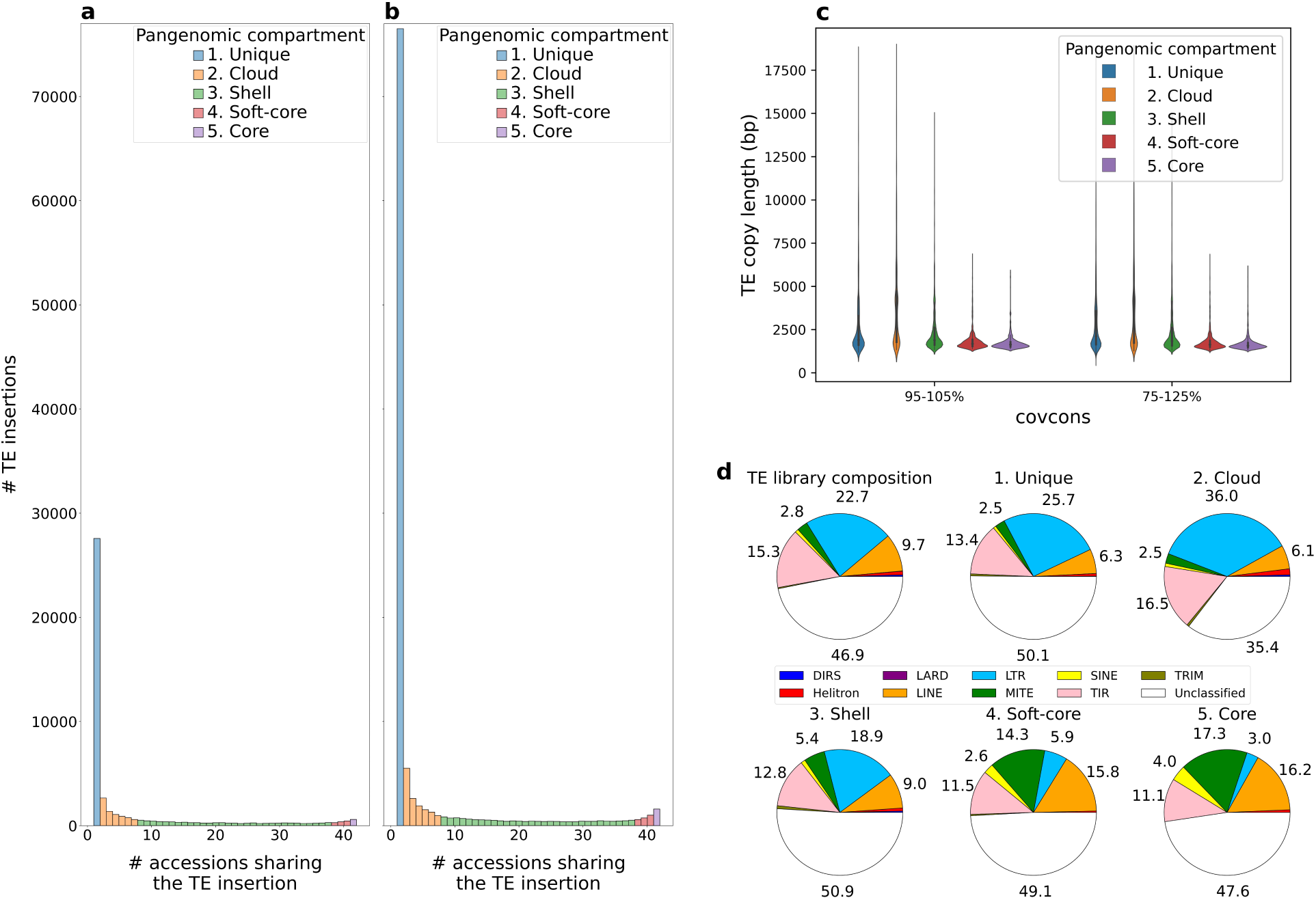
**a, b** Histograms of the number of TE insertions according to the number of accessions sharing each insertion, and the percent coverage between the TE copy and its consensus (parameter covcons). **a** covcons = 95-105% **b** covcons = 75-125% **c** Distribution of TE copy lengths plus their flanking regions according to their pangenomic compartment. **d** Composition of the TE library built *de novo* from Bd21 genome (1,995 sequences) and proportions of TE insertion orders per pangenomic compartment. Only proportion labels greater than 2% are displayed.

We built a clustermap showing the TE copy presence or absence for each accession (**Fig. 3a, 3b)**. Gordon *et al*. estimated three genetic clusters based on >3 million SNPs (Gordon *et al*., 2017): (i) EDF+ with extremely delayed flowering phenotype, (ii) T+ from Eastern, predominantly Turkish, and (iii) S+ from Western, predominantly Spanish. As a validation, we observed that relationships between accessions based on whole-genome SNPs is similar to those computed from shared TE insertions (**Fig. 3a, 3b)**. However, the TE insertion-based tree (**Fig. 3b**) fits even better with the gene-based tree from the Gordon *et al*. study (Gordon *et al*., 2017) (tree not shown); we observed the same two distinct sub-clusters in the cluster T+. Bd29-1 is not clustered as expected and presents few TE insertions contrary to other accessions from its genetic cluster. We think that it does not cluster with its genetic cluster due to its poor assembly quality (the lowest assembled genome size which is 253 Mbp and a low contig N50 of 13 kbp). In order to validate this hypothesis, we took a detailed look at the number of unique TE insertions according to assembly quality metrics **(SF 1, SF 2)**. We see that distribution is not uniform among accessions and does not seem to be explained by genetic clusters **(SF 1**). The number of unique insertions seems to decrease with assembly quality **(SF 2)**. Among the assembly quality metrics, the contig N50 best explains the number of unique TE insertions **(SF 2**) (linear regression with contig N50 as regressor and the number of unique TE insertions as explained variable R² = 0.191, *p*-value = 1.190e-03). We see that the best assembled genomes contain on average fewer unique TE copies than the others **(SF 1)** probably because their copies are well detected and therefore easier to put into a clique.

**Figure 3:**
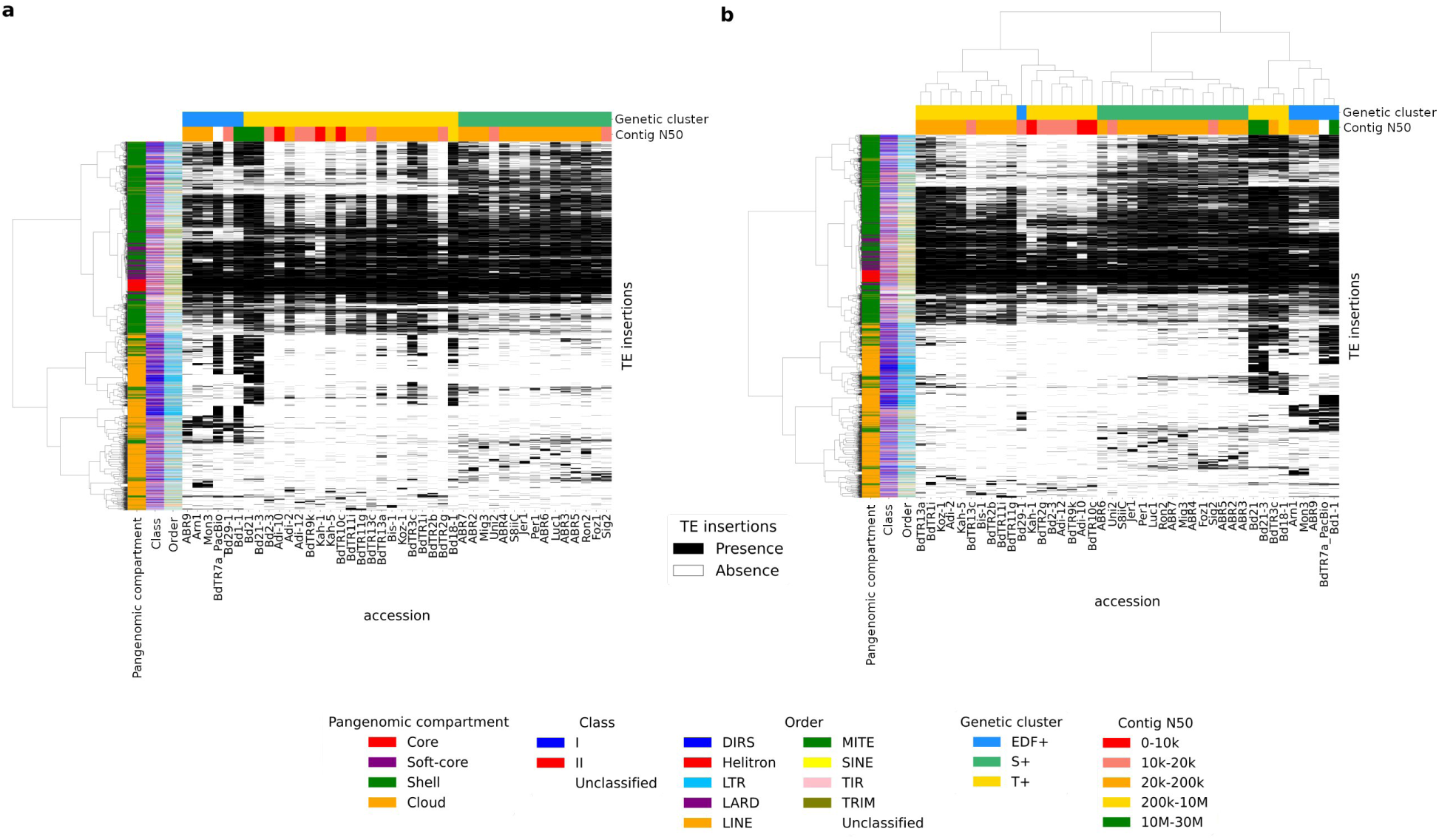
Clustermaps showing TE insertion presence/absence for each accession. **a** Accessions are vertically ordered according to a whole-genome SNP genetic tree (Gordon *et al*., 2017, Supplementary Figure 4.a). **b** Accessions are vertically ordered by their shared TE insertion pattern, after clustering on presence/absence in accessions. The metric used is the Dice distance with Ward’s algorithm.

Unique, cloud and shell TE insertions concern mainly LTR families and the proportion of LTR families decreases as insertion becomes increasingly shared (**Fig. 2d**). Stritt *et al*. showed that the western ancestral population represented by the S+ cluster lost retrotransposons because of several bottlenecks (Stritt *et al*., 2018). Supporting this hypothesis, we observe that the cluster S+ shows low intra-diversity contrary to the other genetic clusters (**Fig. 3a**). For soft-core and core TE insertions, MITE, LINE and SINE proportions increase when insertion is more broadly shared among accessions (**Fig. 2d**). A study on wheat has already shown high sequence conservation of MITEs (Yaakov *et al*., 2013). Horvath *et al*. showed for 320 *B. distachyon* accessions, that retrotransposons are strongly correlated with the demographic structure whereas DNA-transposons are weakly correlated (Horvath *et al*., 2023). This corroborates our observations and validates panREPET method : core and soft-core TE insertions are mainly composed of DNA-transposons, particularly MITE families, while shell, cloud and unique insertions are mainly composed of LTR families and hence shape the intra-specific diversity (**Fig. 2d**).

### Benchmarking panREPET

We compared panREPET against several alternative approaches utilized in recent TE pangenomic studies. Initially, we compared against TEMP (Zhuang *et al*., 2014), which employs paired-end read mapping and is the tool used in the study by Stritt *et al*. for the analysis of TE polymorphisms (Stritt *et al*., 2018). Additionally, a comparison was made with two other approaches that detect structural variations (or SVs) among whole-genome assemblies: Minigraph (Li *et al*., 2020) and GraffiTE (Groza *et al*., 2024).

Stritt *et al*. used the TEMP tool on the 53 *B. distachyon* genomes. They detected 1,889 TE absence polymorphisms (TAPs) which are TEs that are found in the reference assembly but missing in at least one other accession. They also detected 3,627 TE insertion polymorphisms (TIPs) which are insertions not shared with the reference genome Bd21 (Tables S2 and S3 from Stritt *et al*., 2018). We ran panREPET with same Stritt *et al*. genomes and same TE reference library (see Materials and Methods – Benchmarking panREPET). We can only compare shared TE insertions based on genomic coordinates on Bd21 reference genome. Here, panREPET extracted TE copies that cover their reference on more than 80%. panREPET detected 44,597 shared TE insertions (or cliques) whose 24,085 TE copies are shared with the reference Bd21 (equivalent of TAPs) and the others are not shared with the reference Bd21 (equivalent of TIPs) **(Supplementary Table 3)**.

About TAPs, we cannot fetch the coverage copy-consensus from TEMP because the exact consensus is often unknown (Tables S2 and S3 from Stritt *et al*., 2018). In order to retrieve the exact consensus: we intersected TAPs from TEMP against the annotation performed by panTEannot and we selected intersections with the same three-letter code from Wicker’s classification (Wicker *et al*., 2007). We retrieved 1,204 TAPs annotated by panTEannot (among the 1,889 TAPs) **(Supplementary Table 3)**. Then, we filtered TAPs by keeping those that intersect TE copies in Bd21 covering their reference on more than 80% according to panTEannot: we selected 396 TAPs among the 1,204 TAPs **(Supplementary Table 3)**. Among this subset of copies covering 80% of their consensus, TEMP tool finds only 1.6% (396/24,085) of shared TE insertions found by panREPET **(Supplementary Table 3)**, we see that panREPET has detected more shared TE insertions. The number of accessions (TEMP) minus the number of accessions (panREPET) per common shared TE insertion is in mean 25 and in median 25 **(Supplementary Table 3)** that means TEMP tends to overestimate while panREPET underestimates.

About TIPs, we cannot compare TIPs results correctly against panREPET because the genomic coordinates from genomes which are not the reference are not available from TEMP results. For instance, we cannot calculate copy and its consensus coverages because TE sequence lengths of genomes other than the reference are unknown. We retrieve only 3.2% of TE insertions detected by panREPET and shared by Bd21 at TIPs positions **(Supplementary Table 3)**. Note that we considered as retrieved insertion, a TE insertion annotated by the same TE consensus between TIPs from TEMP and panREPET results. This small proportion is expected as TIPs are insertions not shared by the reference Bd21.

Second, we compared shared TE insertions detected by panREPET with those that can be detected as structural variants (SVs) by pangenome graph-methods. We used the sequence-to-graph mapper and graph constructor Minigraph (Li *et al*., 2020): we detected SVs among the 42 *B. distachyon* genomes in graph form (see Materials and Methods - Benchmarking panREPET). SVs are represented as bubbles where each accession has an allele. The questions we asked are: How many TE copies from panREPET intersect an allele from Minigraph? Are they the same accessions between the TE insertion related to the TE copy and the allele in the bubble? We did not consider the core TE copies from panREPET as our aim is to compare against structural variants that are copies shared by a subset of accessions.

Minigraph heavily depends on the first genomes introduced as its graph construction consists in adding genomes incrementally. Hence, we sorted the input genomes in descending order of assembly quality (we used the assembled genome size after removing ‘N’ occurrences which is a a compromise between N50 contigs and N50 scaffolds, see Results - panTEannot). We submitted as the first genome BdTR7a in PacBio because it is the only genome in long read among dataset. Minigraph was configured by default including only SVs affecting 50 bp of sequence or more. Minigraph detected 181,965 segments, 90,705 alleles, 44,163 bubbles and 3 alleles/bubble (mean and median). Alleles are a combination of segments. Alleles have a mean of 5 kbp and a median of 122 bp, whereas TE copies present less variance with a mean of 1.7 kbp and a median of 979 bp **(SF 5a-b)**.

We considered two cases, the first being when the allele is totally included in TE copy (case 1) and the second when the TE copy is totally included in allele (case 2). Case 1 highlights intra-copy polymorphism and case 2 TE copy polymorphisms (or TE insertions). Regarding the 231,510 TE copies from panREPET detected among 42 accessions (for covcons=95-105%), 43.5% of them overlap an allele from Minigraph with at least 10% coverage between the allele and the TE copy, (**Table 2**). Considering only BdTR7a and the reference genome Bd21, the proportion of TE copies matching an allele is higher (53.7% and 54.8% respectively), which is expected, since the graph construction with Minigraph depends on the first introduced genomes (it increments its graph by adding genomes) (**Table 2**). About the 56% remaining of TEs which do not overlap an allele from Minigraph, they exhibit a median of 831 bp and an average of 1.3 kbp, whereas TE copies overlapping an alleles are slightly larger with a median of 1.1 kbp and a mean of 2.1 kbp **(SF 5a)**. Largest alleles overlap a TE copy with a median of 210 bp and a mean of 9.5 kbp, whereas alleles not overlapping a TE copy have a median of 100 bp and a mean of 2.8 kbp **(SF 5b)**.

**Table 2:** Intersection results between TE insertions detected by panREPET against alleles detected by Minigraph. TE copies covering 95-105% of their consensus are in black, and coverages between 75-125% are in blue.

|  |  | All intersections |  |  | Case 1: The allele is included in TE copy |  | Case 2: The TE copy is included in allele |  |
| --- | --- | --- | --- | --- | --- | --- | --- | --- |
| Minimum TE copy-Allele coverage (%) |  | 10 | 50 | 100 | 50 | 100 | 50 | 100 |
| Proportion of TE copies matching an allele (%) | All accessions | 43.5<br>43.1 | 41.1<br>40.6 | 36.1<br>35.6 | 19.2<br>19.3 | 13.1<br>13.3 | 26.8<br>26.1 | 23<br>22.3 |
|  | BdTR7a | 53.7 | 51 | 45.4 |  |  |  |  |
|  | Bd21 | 54.8 | 51.9 | 45.5 |  |  |  |  |
| Number of TE copies / allele | mean | 7.4 | 7.2 | 7.1 | 7.5 | 8.9 | 6.4 | 6.4 |
|  | median | 3 | 3 | 3 | 3 | 4 | 3 | 3 |
| Number of alleles / TE copy | mean | 1.2 | 1.1 | 1.1 | 1.2 | 1.3 | 1 | 1 |
|  | median | 1 | 1 | 1 | 1 | 1 | 1 | 1 |

To assess the fragmentation in the intersections alleles from Minigraph against TE copies from panREPET, we fetched the number of TE copies intersecting an allele in average and the number of alleles intersecting a TE copy in average (**Table 2**). As alleles are on average larger than TE copies, it was expected that the average number of TE copies per allele would be greater than 1, and we found 7 in mean and 3 in median (**Table 2**). It corresponds to large SVs containing several TE copies. We also observed that there are about 1.1 alleles per TE copy on average (**Table 2**), which means that some TE copies are fragmented into several alleles **(SF 6)**.

To assess the detection differences between panREPET and Minigraph, we calculated for each TE copy-allele intersections covering over 100% (on the allele for the case1, on the TE for case2), the difference: number of accessions sharing the TE copy minus the number of accessions sharing the associated allele. **SF 5c-j** are histograms showing the distribution of those differences. On the left the differences mean that panREPET underestimates compared to Minigraph the number of accessions sharing the TE copy and on the right, it overestimates. We calculated then the proportion of common accessions (**SF 5c-d**). The more the difference is around 0 and the proportion is close to 100%, the more the shared TE insertion detected by panREPET fits with SVs detected by Minigraph. There are fewer cases where the allele is included in the TE copy (case 1, **SF 5c**). In this case, panREPET tends to underestimate the number of accessions and Minigraph to overestimate **(SF 5c)**. This shows that when the TE copy is incompletely detected, Minigraph loses specificity. This concerns a large part of large TE copies (more than 4kbp) (case 1, **SF 5g**) which corresponds in majority to unique and cloud TE copies (**Fig. 2c, SF 5e)**. That means that for the larger TE copy, Minigraph loses specificity **(SF 6)**. We observe more case 2 when the TE copy is in allele, both tools overestimate or underestimate the number of accessions **(SF 5d)**. This shows that Minigraph does not detect variations at the TE scale, but at the SV scale which is a larger scale. Hence, an approach detecting SVs firstly then annotating SVs in TEs is limited. By adding more partial copies with panREPET (i.e. TE copies covering 75-125% of their consensus), observations remain unchanged.

Third, we tested GraffiTE. As for TEMP, we were just able to compare genomic coordinates based on the reference genome Bd21. Indeed, GraffiTE compares each alternative genome against the reference without giving TE coordinates on alternative genomes. GraffiTE detected 47,236 SVs whose 111,860 TE copies are deletions and 48,241 TE copies are insertions among all alternative accessions **(Supplementary Table 4)**. A deletion is here a TE copy absent in an alternative genome but present in the reference genome. A deletion corresponds to a shared TE insertion in panREPET which is also shared by Bd21 but absent in at least one genome. An insertion is a TE copy present in at least one alternative genome but absent in the reference Bd21. A SV detected by GraffiTE can contain several detected TEs (here around 2-3 TE copies per SV, **Supplementary Table 4**).

For comparing against panREPET, we only extracted TE copies from GraffiTE covering their consensus between 95 and 105%. Among deletions, that corresponds to 6,487 TE copies on 111,861 total TE copies from all accessions (5.7%) and corresponds also to 3,166 shared TE insertions with reference genome Bd21 **(Supplementary Table 4)**. Among insertions, that corresponds to 1,169 TE copies are insertions among 48,241 TE copies insertions (2.4%, **Supplementary Table 4**). After intersecting deletions and insertions based on the reference Bd21 genome from GraffiTE against Bd21 shared TE copies from panREPET, we considered as a retrieved TE copy, a copy annotated by the same consensus. We do not consider core TE insertions from panREPET as GraffiTE does not aim to detect core TE insertions. Among deletions from GraffiTE, panREPET retrieved 1,802/3,166 (56%) (**Figure 4, Supplementary Table 4)**. Among insertions from GraffiTE, panREPET retrieved only 23 copies among 1,169 (1.9%), which is expected as insertions are not shared by the reference Bd21 **(Supplementary Table 4)**. Among Bd21 shared TE copies from panREPET, GraffiTE retrieved 1,802/9,474 (19%) (**Figure 4, Supplementary Table 4)**.

**Figure 4:**
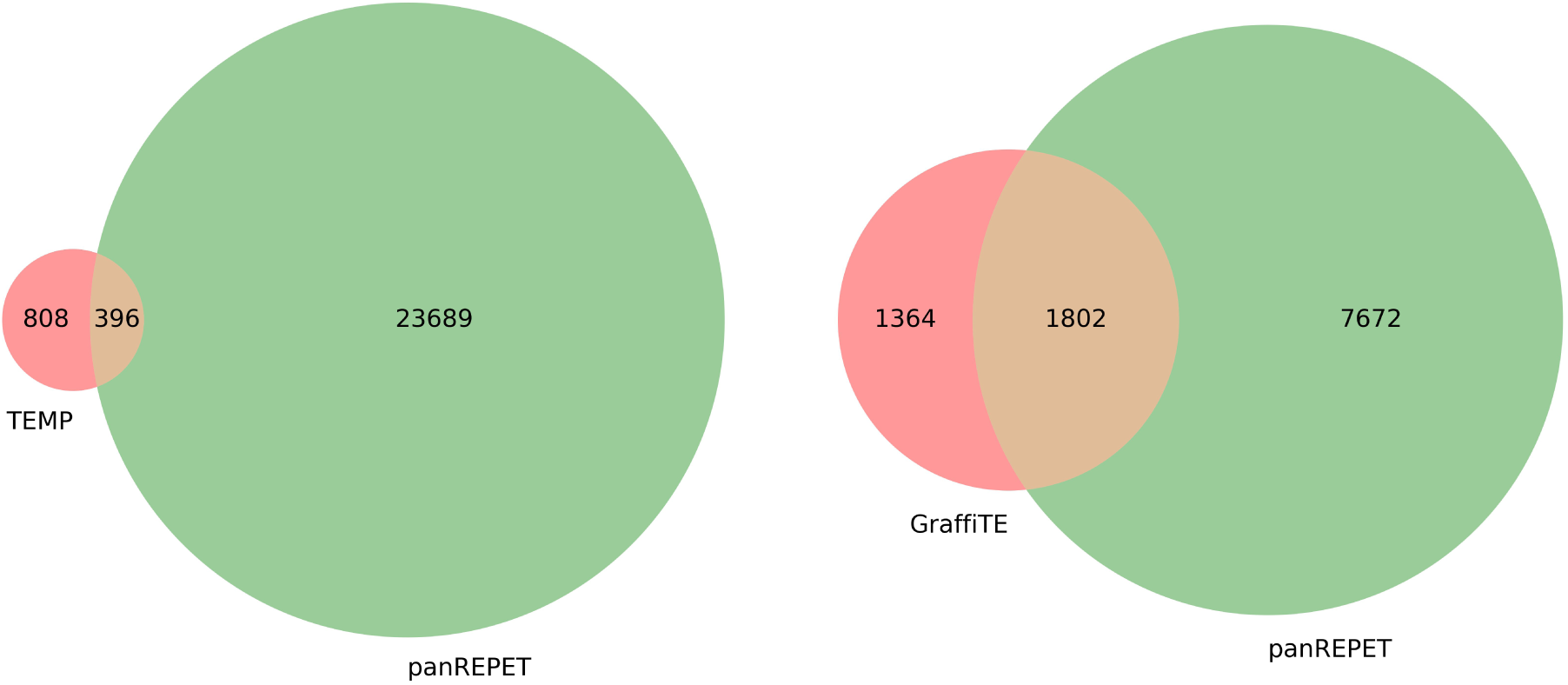
Venn diagrams showing shared TE insertions with the reference genome found in common or not between the two compared tools (left against TEMP tool and right against GraffiTE tool). We compared by selecting TE copies covering their reference on more than 80% for TEMP and 95% for GraffiTE.

GraffiTE detected in mean and in median 1 accession per SV (deletion or insertion), whereas panREPET detected in mean 19 and in median 18 accessions sharing a TE copy. GraffiTE tends to overestimate the number of individuals sharing a TE insertion (number of accessions from GraffiTE minus number of accessions from panREPET per common shared TE insertions are in mean 26 and in median 31, **Supplementary Table 4**). GraffiTE also detected better cloud and unique TE copies (30% and 32% respectively, **Supplementary Table 4**) than shell and soft-core shared TE insertions from panREPET (16% and 5.9% respectively, **Supplementary Table 4**). About sequence lengths, GraffiTE detected better large TE copies among TE copies from panREPET : copies retrieved have a mean of 3,3 kbp and 3,0 kbp in median whereas copies not retrieved have a mean of 3,2 kbp and 2,4 kbp in median **(Supplementary Table 4**).

### Core TE insertions

Within the pangenome, about 10% of core TE insertions intersect at 80% a feature of a gene (**Table 3**). They mainly intersect exons (**Fig. 5a**). On the Bd21 genome and among pangenomic TE compartments, between 1.3% and 4% of the Transcription Factor Binding Site (TFBS) in promoter regions (−500 bp ∼ +100 bp of transcription start site (TSS)) from the PlantRegMap database (Tian *et al*., 2019) intersect a TE (**Table 3**). To test whether these proportions significant, we shuffled the 17,589 predicted TFBS in promoter regions on the Bd21 genome thanks to bedtools shuffle (Bioinformatics, 2024) and counted the number of intersections against core TE insertions with 100% coverage on the TFBS. The null hypothesis is that there is a statistical association between core TE insertions and the shuffled TFBS only by chance because of their numbers. Repeating this procedure 100 times, we observed that shuffled TFBS do not intersect the core TE insertions (Chi-Square goodness of fit = 714, *p*-value = 0.0, degrees of freedom = 99). We can conclude that there is a statistically significant association between core TE insertions and TFBS in promoter regions, suggesting that some of the core TEs result from co-optation of a TE-TFBS (Baud *et al*., 2019). Concerning the 144,761 genome-wide predicted TFBS (i.e., all TFBS including those not in known promoter regions), there is also no statistical association by chance between core TE insertion and all TFBS (Chi-Square goodness of fit = 8,042, *p*-value = 0.0, degrees of freedom = 99). In the following, we will work only with TFBS predicted in promoters. We observe that core TEs with TFBS are mostly MITEs (**Fig. 5d-e**) as also shown on maize where domesticated MITEs carrying TFBS appear to be involved in husk tissue-specificity (Fagny *et al*., 2021).

**Figure 5:**
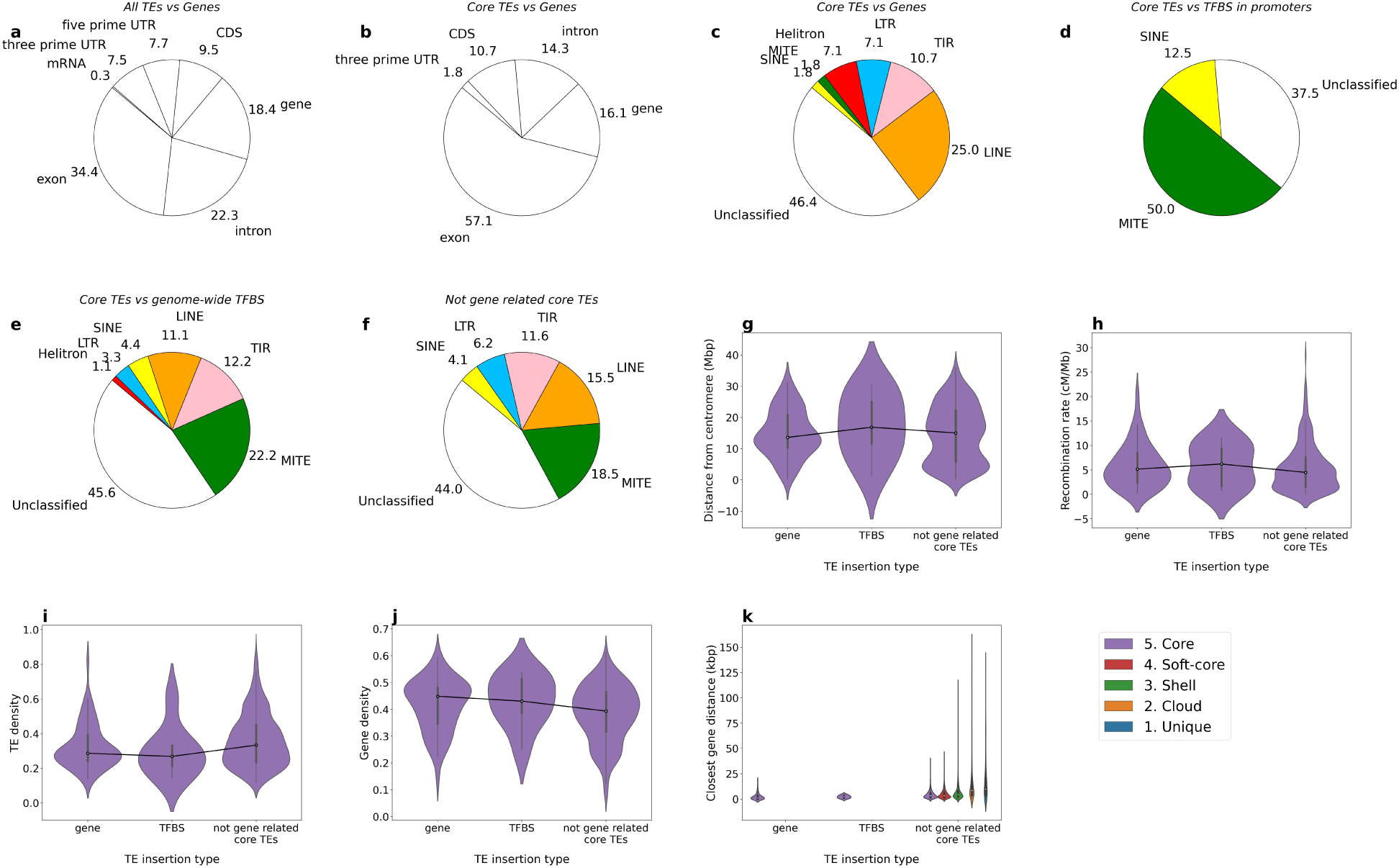
Analysis of core TE insertions on the Bd21 genome. **a -** Gene features that all TE copies from Bd21 genome intersect by at least 80%. **b -** Gene features that the core TE insertions from Bd21 genome intersect by at least 80%. **a-b -** Only the gene feature most covered by the TE is selected. **c-f -** Order of TE family proportions that the core TE insertions from Bd21 genome intersect. **c -** Core intersecting a gene. **d** - Core intersecting a TFBS in promoter regions. **e -** Core intersecting a genome-wide TFBS. **f -** *Not gene related core TE* insertions (by considering TFBS in promoter regions). **g-k** - Parameter distributions according to core TE insertions that intersect a gene, those that intersect a TFBS in promoter regions and *not gene related core TE* insertions. **g** - Distance distribution from the centromere (Mbp). **h** - Recombination rate (cM/Mb). **i** - TE density. **j** - Gene density. **k -** Closest gene distance (kbp).

**Table 3:**
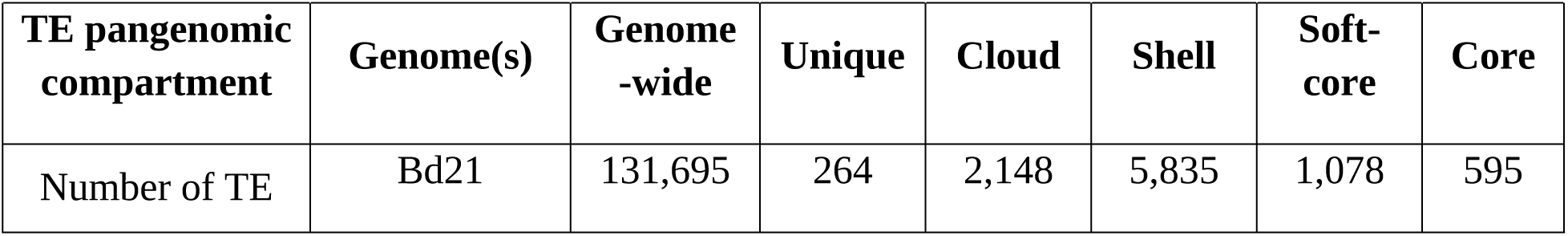
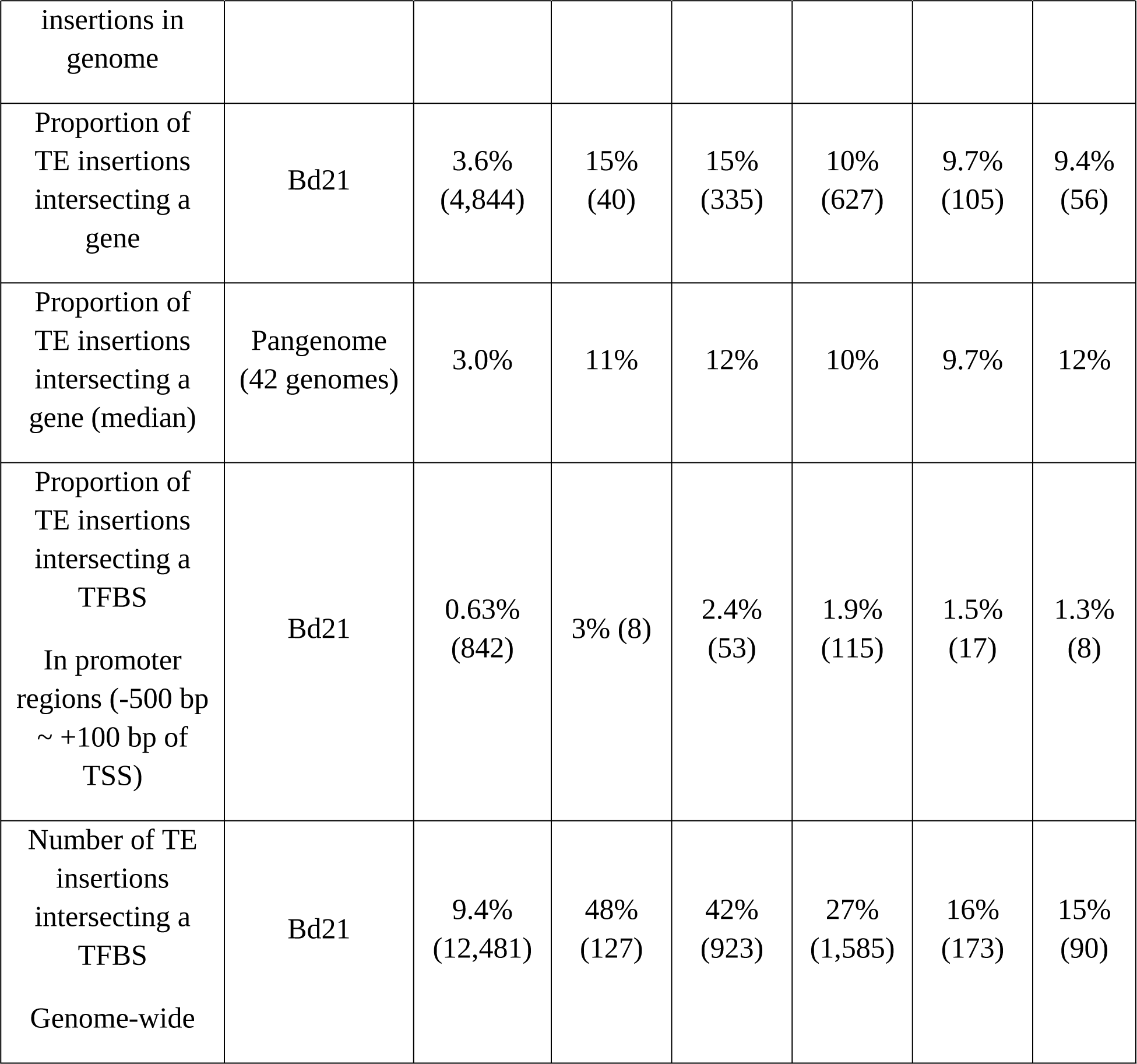
Number of TE insertions that intersect genes (covering at least 80% of a gene feature) or TFBS (covering 100% of the TFBS) in terms of TE pangenomic compartments. TFBS: Transcription Factor Binding Site. TSS: Transcription Start Site.

Concerning the remaining core TEs not overlapping genes or TFBS (534/595 core TE insertions) called hereafter *not gene related core TEs* (**Table 3**), we wondered about their conservation at the species level. First, we hypothesize that TEs may be domesticated thanks to their role as essential structural components for centromeres (Casola *et al*., 2008 ; Wong *et al*., 2004) and for maintaining centromeric and telomeric stability or heterochromatic silencing (Gao *et al*., 2005). To determine whether the *not gene related core TE*s have been conserved thanks to these roles, we plotted their positions on chromosomes **(SF 4)** and determined their distance from the centromere, the recombination rates of the region of insertion (cM/Mb), the TE and gene density, and their distance to the closest gene (**Fig. 5g-k).** We performed an ANOVA to compare these distributions between *not gene related core TE*s and the other core TE insertions. We observed that only one *not gene related core TE* is located at the centromere, and these core TE insertions seems more closer to the centromere than core TE insertions intersecting a gene (**Fig. 5g, ANOVA *p*-value=3.648e-03)**. TE insertions are also more likely to settle in non-recombining regions by fixation (Bartolomé et al., 2004) as they may have been fixed by genetic drift (Jurka *et al*., 2011 ; Almojil *et al*., 2021). *Not gene related core TE*s are not more frequent in low-recombination regions than the other core TEs (**Fig. 5h, ANOVA *p*-value=0.4576).** Heterochromatic regions are also less subject to recombination (Straub *et al*., 2003), and TEs tend to accumulate there (Charlesworth *et al*., 1994 ; Bartolomé *et al*., 2002 ; Capy *et al*., 2021). To infer the chromatin state of the genomic regions, we computed respectively TE and gene density per 1Mbp intervals among chromosomes (**Figs. 5i-j**). We observe that *not gene related core TE*s are not preferentially located in heterochromatin regions (**Figs. 5i-j, ANOVA *p*-values=0.1395 and 0.0516 for TE density and gene density respectively)**. However, these TE insertions are further from the closest gene than the other core TEs (**Fig. 5k, ANOVA *p*-value=4.751e-03)**.

Regions under negative selection and low recombination rates do not allow efficient elimination of transposable elements (the Hill-Robertson effect) (Bartolomé *et al*., 2002 ; Charlesworth *et al*., 2021). Consequently, we observe on **Fig. 5k** that *not gene related unique and cloud TE* insertions present a greater distance to the closest gene compared to the *not gene related core and soft-core TE* insertions **(T-test: T-statistic=-19.96, *p*-value=2.54e-84)**. We can then hypothesise that *not gene related core TE* insertions are conserved by the effect of a nearby gene under positive selection.

### TE insertion dates

Age of TE insertion events can highlight their evolutionary dynamics over time. Combining genetic distances between accessions with our analysis of TEs in the pangenome allowed us to date TE insertions more accurately than current methods since we take into consideration divergence between accessions. In addition, we consider all types of TE copies and not only the LTR copies contrary to others methods (SanMiguel *et al*., 1998 ; Vitte *et al*., 2007 ; Daron *et al*., 2014 ; Stritt *et al*., 2020). Some other methods date TE copies using percent identity with the consensus as a proxy of age. The more the copy diverges from the consensus, the older it would be (Britten *et al*., 1988; Maumus *et al*., 2016). For all TE copies detected among accessions, we represented the distribution of TE copy-consensus identity percentage, according to the number of accessions sharing the TE insertion **(SF 3)**. As expected, core TE insertions **(SF 3)** show low identities with their consensus sequence (<65%), suggesting ancient and conserved copies.

This approach is however limited as it depends on consensus from the reference TE library. We propose a new approach for dating TE insertions based on whole-genome SNPs that takes into account species-wide evolution. TE insertions shared between accessions are insertions that preceded the split of lineages . TE insertion age can be then inferred as the longest whole-genome SNP distance between each pair of accessions sharing the TE insertion. We observe that the more an insertion is shared, the greater the genetic distance based on SNPs (**Fig. 6a**). The observed plateau is the maximum SNP distance (0.335 subs/site) among accessions, corresponding to distance between the two most distant accessions Arn1 and Ron2 (see Gordon *et al*., 2017, Supplementary Figure 4.a). Minadakis *et al*. estimated the genetic cluster divergences based on a multispecies coalescent approach (Minadakis *et al*., 2023): S+, T+ and EDF+ genetic clusters split at least 45 kya ago. More recently, S+ and T+ genetic clusters split 23 kya ago. For converting substitution per site into kya, we assumed that the maximum genetic distance based on whole-genome SNPs among S+ and T+ genetic clusters (0.1549 subs/site, between Bd21 and Sig2) corresponds to 23 kya.

**Figure 6:**
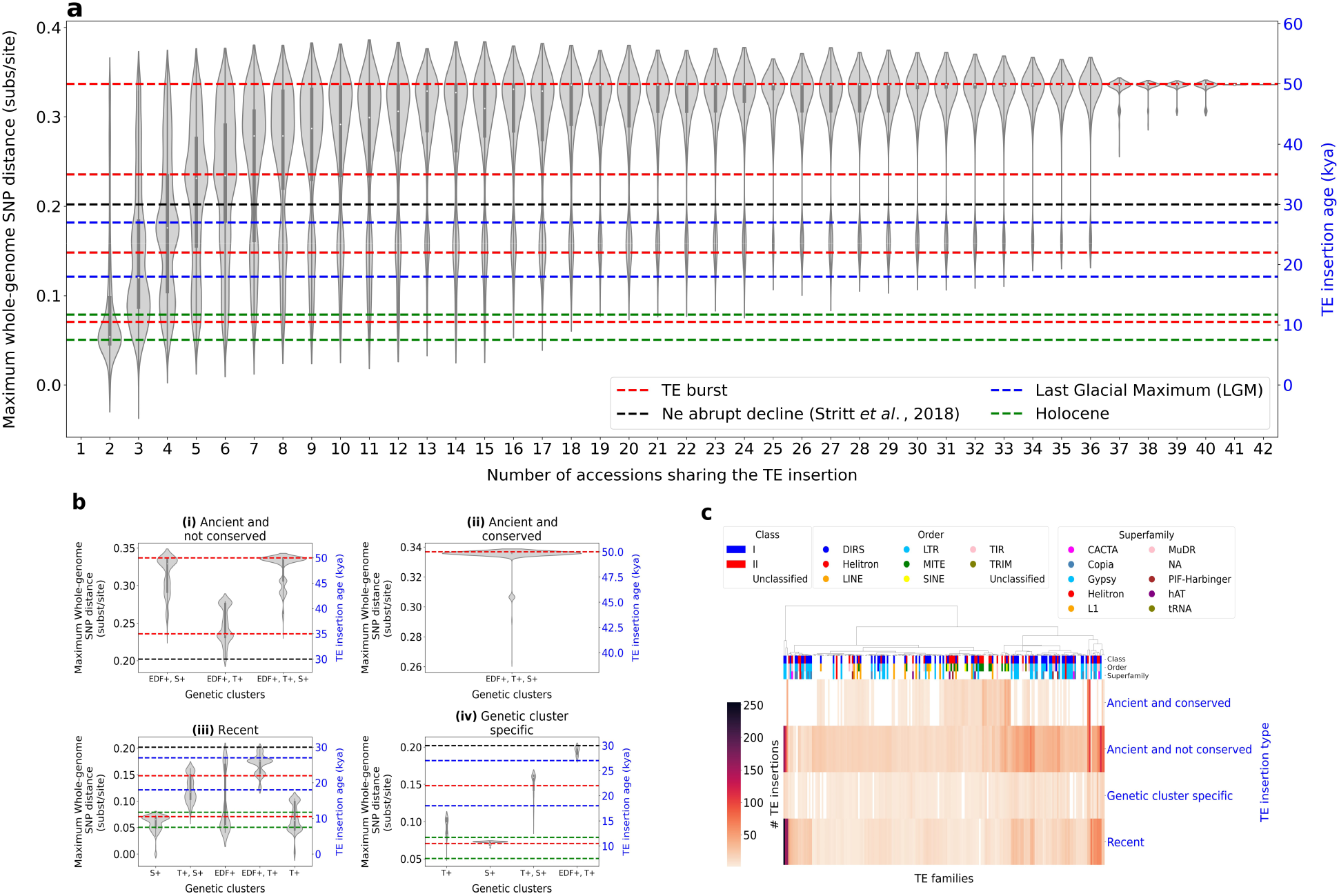
a - Boxplots showing age distribution of TE insertions according to the number of accessions sharing them. The age is estimated by the maximum pairwise whole-genome SNP distance between accessions within the TE clique. **b -** Boxplots showing age distribution of TE insertions according to genetic clusters to which the accessions sharing the insertion belong. We split Fig 5a. into four TE insertion types: (i) the ancient and not conserved TE insertions; (ii) The ancient and conserved TE insertions; (iii) The recent TE insertions and (iv) the TE insertions that occurred after the cluster divergence. **c -** Clustermap showing the abundance of TE families according to the TE insertion types. The metric used is the Euclidean distance with ward algorithm. We selected families having more than 20 copies for readability.

**Fig. 6a** illustrates four types of TE insertion events: (i) The ancient and not conserved TE insertions are shared by few to many accessions (2-36 accessions) and at least two accessions are genetically distant on the whole-genome-SNP genetic tree (i.e., with a genetic distance greater than 0.20 subs/site corresponding to 30 kya). We counted 9,824 among the 18,513 shared TE insertions. However, the accessions that most share this type of insertion are Bd1-1, Bd18-1, Bd21, Bd21-3, BdTR3c and BdTR7a. Note that they have high quality assembly **(SF 1, Fig. 3**), and it is more likely that they share these insertions because the corresponding regions are better assembled than in other genomes. This type of TE insertion is numerous and Horvath *et al*. showed that TEs in *B. distachyon* are dominated by purifying selection (Horvath *et al*., 2023), which corroborates our observation. (ii) The ancient and conserved TE insertions are shared by almost all accessions (37- 42 accessions) suggesting a pre-speciation insertion. We counted 2,296 among the 18,513 shared TE insertions. (iii) The recent TE insertions are shared by only few accessions (2-6 accessions) which are close on the SNP genetic tree (genetic distance smaller than 0.20 subs/site or 30 kya). We counted 4,890 among the 18,513 shared TE insertions. In this case, the common ancestor underwent insertion recently. They mainly concern the couple Arn1 and Mon3 from Spain. They specifically share a mixed ancestry (Minadakis *et al*., 2023), that explains why they share so many specific TE insertions. (iv) The TE insertions that occurred after the divergence into intra-specific genetic clusters are only shared by accessions from an intra-specific clade. We counted 1,503 among the 18,513 shared TE insertions.

We dated four major transposition events in *B. distachyon*. A first major transposition event, at least at 45 kya, that could correspond to pre-speciation TE insertions or TE insertions occuring during the split into the three lineages. A second major transposition event at 35 kya that involves TE mostly shared by EDF+ and T+ (**Fig. 6b**). That could suggest that the S+ genetic cluster underwent genetic drift around 35 kya with the hypothesis that the few S+ individuals surviving the bottleneck have their copy eliminated by genetic drift or selection. A third major transposition event at 22 kya during the Last Glacial Maximum (LGM), a period conducive to thermal stress. Note in **Fig. 6a** that we observe a gap of TE insertions around 30 kya that may be explained by the abrupt decline in population sizes undergone by the species, followed by population expansion in the very recent past (Stritt *et al*., 2018 ; Minadakis *et al*., 2023). These population expansions may have triggered TE transposition around 22 kya. Finally around 10 kya, there is a burst of transposition occurring during the Holocene (11.7-7.5 kya) which is the beginning of deglaciation in Europe and the relocation of species to new environments (Minadakis *et al*., 2023). Among the recent transpositions, we observed a burst of transposition in S+ at 11.7 kya (**Fig. 6b**). S+ may have experimented bursts of transposition during its recolonization of Europe and the Middle East.

Identifying active families that are present at very high copy number allows us to understand the impact of TEs on plant genome evolution (Feschotte *et al*., 2002). A way to highlight families that have transposed during these major transposition events, is to represent the abundance of TE families according to their insertion type (**Fig. 6c**). We see that the oldest conserved transpositions, that occurred at least 45 kya, involve in majority MITE, LINE, TIR and SINE families. LTR families are however either recent or ancient and not conserved, suggesting an active turnover: the two LTR families (on the left in **Fig. 6c**) correspond to Ty3/Gypsy. Stritt *et al*. suggested that recent TE transpositions involve centromeric Ty3/Gypsy superfamily and Ty1/Copia superfamily (Stritt *et al*., 2020).

### Factors affecting the transposition dynamics of TE families

We investigated environmental factors that may explain TE mobilization by using geo-climatic data. We are interested here in local factors which could explain recent TE transposition. In order to highlight the differential dynamics of recent TE families between accessions, we clustered the accessions according to the number of recent TE family copies (**Fig. 7**). As a first proxy, recent TE insertion can be identified as those that are unique, i.e. occurring within only one accession. We distinguished unique TE insertions with an identity TE copy-consensus greater than 95% (95% is a cut-off already used to highlight young subfamilies (Kapitonov *et al*., 2003) (**Figs. 7a, 7a’)** and the other ones (**Figs. 7b, 7b’).** Another way to identify recent transposition events is by selecting only TE insertions occurring after the Holocene (i.e., age < 7.5 kya). Indeed, we chose this threshold in particular because *B. distachyon* moved southward during the last glacial period and recolonized Europe and the Middle East within the last five thousand years (Niche modeling analysis performed by Minadakis *et al*., 2023) (**Figs. 7c, 7c’)**.

**Figure 7:**
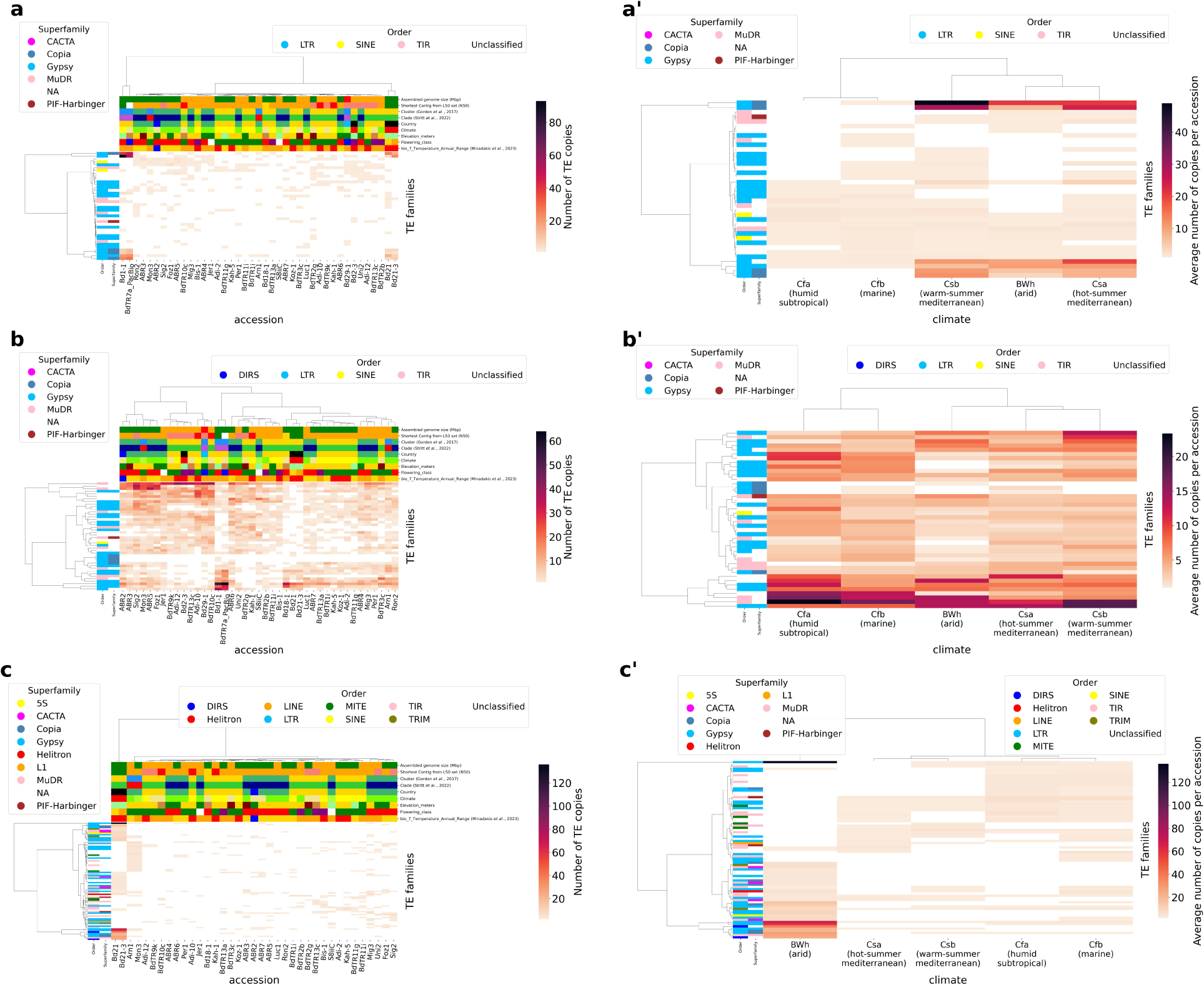
a-c- Clustermaps showing the abundance of recent TE insertions per TE families and per accession. On the x-axis and on the y-axis, accessions and TE families are respectively clustered according to their similarity in their pattern of TE family dynamics. Color labels of columns are detailed in **Supplementary Table 2**. We used the Euclidean distance with the ward linkage method. The percent coverage between TE copy and its consensus is 95-105%. **a-b** - Unique TE copies. Only TE families with at least 10 insertions for at least one individual have been selected. **a** - TE copies with an identity greater than 95% with their consensus. **b** - TE copies with an identity lower than 95% with their consensus. **c** - TE copies that transposed after the Holocene (age < 7.5 kya). **a’-c’ -** Each clustermap corresponds to its counterpart, the only differences are that the climate is on the x-axis and the clustermap shows the average number of TE copies per accession in order to weight the unequal distribution of accessions in the different climate classes (Arid BWh: 3 accessions, Humid subtropical Cfa: 3, Marine Cfb: 14, Hot-summer mediterraneen Csa: 13, Warm-summer mediterraneen Csb: 9).

**Table 3:** Search on Bd21 genome for genome-wide predicted TFBS intersecting recent and over-represented TE insertions (covering 100% of the TFBS).

| Figures | 6a, 6a' | 6b, 6b' | 6c, 6c' |
| --- | --- | --- | --- |
| **Recent TE insertion criteria** | Unique TE insertions<br>Copy-consensus percent identity > 95% | Unique TE insertions<br>Copy-consensus percent identity <= 95% | After the Holocene (age < 7.5 kya) |
| **Supernumerary TE families criteria** | TE families with more than 10 copies | TE families with more than 10 copies | TE families with more than 2 copies |
| **TE families intersecting a TFBS** | MCL2 LTR Gypsy: 12/12 copies<br>MCL14 LTR Copia Angela: 19/19 copies<br>MCL42 LTR Copia: 4/4 copies | MCL2 LTR Gypsy: 18/21 copies<br>MCL4 LTR Gypsy: 9/12 copies | MCL2 LTR Gypsy: 28/28 copies<br>MCL4 LTR Gypsy: 22/22 copies<br>MCL3 TIR MuDR: 13/13 copies<br>MCL14 LTR Copia: 11/11 copies<br>MCL68 LTR Copia: 10/10 copies |
| **Transcription Factors inferred from rice** | MCL2: ('AP2 domain containing protein, expressed', 7), ('TCP-domain protein, putative, expressed', 5), ('OsMADS56 - MADS-box family gene with MIKCC type-box, expressed', 4), ('OsMADS4 - MADS-box family gene with MIKCC type-box, expressed', 4), ('tesmin/TSO1-like CXC domain containing protein, expressed', 3), ('TCP family transcription factor, putative, expressed', 2), ('MYB family transcription factor, putative, expressed', 2)<br>MCL14: ('dof zinc | MCL2: ('AP2 domain containing protein, expressed', 7), (nan, 7), ('TCP-domain protein, putative, expressed', 5), ('TCP family transcription factor, putative, expressed', 4), ('OsMADS56 - MADS-box family gene with MIKCC type-box, expressed', 4), ('tesmin/TSO1-like CXC domain containing protein, expressed', 3), ('MYB family transcription factor, putative, expressed', 2), ('OsMADS4 - MADS-box family gene with MIKCC type-box, expressed', 1) | MCL2: ('AP2 domain containing protein, expressed', 13), ('OsMADS56 - MADS-box family gene with MIKCC type-box, expressed', 7), ('DUF260 domain containing protein, putative, expressed', 6), ('TCP family transcription factor, putative, expressed', 6), ('MYB family transcription factor, putative, expressed', 4), ('tesmin/TSO1-like CXC domain containing protein, expressed', 4), ('TCP-domain protein, putative, expressed', 4), ('dof |
|  | finger domain containing protein, putative, expressed', 15), ('OsMADS8 - MADS-box family gene with MIKCC type-box, expressed', 7)<br>MCL42: ('Myb-like DNA-binding domain containing protein, putative, expressed', 3), ('OsMADS3 - MADS-box family gene with MIKCC type-box, expressed', 1) | MCL4: ('AP2 domain containing protein, expressed', 12), ('TCP family transcription factor, putative, expressed', 7), ('TCP-domain protein, putative, expressed', 6), ('DUF260 domain containing protein, putative, expressed', 2), ('OsMADS56 - MADS-box family gene with MIKCC type-box, expressed', 1) | zinc finger domain containing protein, putative, expressed', 3), ('single myb histone, putative, expressed', 2), ('OsMADS4 - MADS-box family gene with MIKCC type-box, expressed', 1)<br>MCL4: ('AP2 domain containing protein, expressed', 13), ('TCP family transcription factor, putative, expressed', 4), ('OsMADS56 - MADS-box family gene with MIKCC type-box, expressed', 4), ('DUF260 domain containing protein, putative, expressed', 3), ('Homeobox domain containing protein, expressed', 3), ('bZIP transcription factor, putative, expressed', 2), ('ethylene-responsive transcription factor, putative, expressed', 1), ('tesmin/TSO1-like CXC domain containing protein, expressed', 1), ('MYB family transcription factor, putative, expressed', 1), ('no apical meristem protein, putative, expressed', |
|  |  |  | 1)<br>MCL3: ('ZOS2-02 - C2H2 zinc finger protein, expressed', 7), ('Homeobox domain containing protein, expressed', 6), ('AP2 domain containing protein, expressed', 3), ('CPuORF1 - conserved peptide uORF-containing transcript, expressed', 2), ('bZIP transcription factor domain containing protein, expressed', 2), ('zinc finger C-x8-C-x5-C-x3-H type family protein, expressed', 1), ('TCP family transcription factor, putative, expressed', 1), ('OsMADS56 - MADS-box family gene with MIKCC type-box, expressed', 1), ('GAGA-binding protein, putative, expressed', 1), ('B3 DNA binding domain containing protein, expressed', 1), ('tesmin/TSO1-like CXC domain containing protein, expressed', 1), ('dehydration-responsive element-binding protein, putative, expressed', 1) |
|  |  |  | MCL14: ('dof zinc finger domain containing protein, putative, expressed', 7), ('OsMADS8 - MADS-box family gene with MIKCC type-box, expressed', 6)<br>MCL68: ('helix-loop-helix DNA-binding domain containing protein, expressed', 10) |

Similar patterns of TE family dynamics among a set of accessions may suggest common determinants for TE activity (e.g., environmental stress). To determine those common determinants, we looked at for each accession: its genetic cluster, the clade, the country, the climate, the elevation in meters and the flowering class. Minadakis *et al*. showed that annual mean precipitation presented the highest number of genes located in significantly associated regions (Minadakis *et al*., 2023), for this reason we also added the annual mean precipitation from their analyse. Unique TE family dynamics does not seem to be explained by genetic clusters but by other factors (**Figs. 7a-b**). Accessions presenting the best N50 at the contig level are clustered (the couples Bd1-1/BdTR7a from Turkey and Bd21/Bd21-3 from Iraq) (**Fig. 7a**). They present fewer unique TE insertions than the other accessions **(SF 1)**, yet the same families cluster together, which is not the case in the other accessions. This may suggest that accessions with low assembly quality show many single copies which are actually noise. The couples (Bd21, Bd21-3) and (Bd1-1, BdTR7a) specifically share Copia and Ty3/Gypsy LTR and MuDR TIR superfamilies (**Fig. 7a**).

We also represented the TE family dynamics directly according to climate class for TE insertions occurring after the Holocene (age < 7.5 kya) (**Fig. 7c’)** to highlight transpositions induced by abiotic conditions. We plotted the average number of TE copies per accession to avoid biases from non-homogeneous distribution of accessions in the different climate classes (Arid Bwh: 3 accessions, Humid subtropical Cfa: 3, Marine Cfb: 14, Hot-summer mediterraneen Csa: 13, Warm-summer mediterraneen Csb: 9). It has been observed that accessions in arid climates exhibit a clear over-representation of unique copies. This observation suggests the transposition activation of specific transposable element (TE) families. Furthermore, it can be inferred that accessions in hot summer and warm summer Mediterranean climates are more similar to each other than to those in humid subtropical and marine climates (**Fig. 7c’)**.

Then we looked for TFBS in TEs potentially explaining a local TE activation. We looked at the intersection of the 144,761 predicted genome-wide TFBS with the supernumerary TE copies (i.e., TE families with more than 10 copies or 2 copies, see **Table 3**) and we only kept TEs intersecting by 100% a TFBS sequence (**Table 3**). We only worked on Bd21 because the analysis required available TFBS data. Among the supernumerary recent TE insertions intersecting a TFBS, Bd21 presents in majority LTR families (**Table 3**). We observed TFBS targeted by OsMADS56, OsMADS4, OsMADS8 and OsMADS3 aMADS-box gene family with an MIKCc type-box. Studies on *Oryza sativa* showed that they are critical for the reproductive organ (Fornara *et al*., 2004). OsMADS8 is required for tapetum development (Chen *et al*., 2018). Its protein degradation by certain ambient air pollutants can lead to pollen sterility (Rezanejad *et al*., 2013) in the petunia.

In addition, the myb-like DNA-binding domain such as for instance in AtMYB002, is involved in drought response in *Arabidopsis thaliana* (Ambawat *et al*., 2013).

## Discussion

### panREPET, a new method to improve TE analysis

Several pipelines exist for detecting TEs in pangenomes. They generally analyse TEs according to a reference genome. Two approaches have tried to find neo-TE-insertions in resequenced genomes: (i) paired-end mapping (PEM) detecting discordant pairs (Sabot *et al*., 2011 ; Kofler *et al*., 2016) and (ii) the split-read method detecting truncated reads corresponding to the junction between the TE insertion and the reference sequence. Unlike the strategy of discordant reads, truncated reads can determine the TE insertion at the base level. However, the coverage is low, and this method is very sensitive to read size (Carpentier *et al*., 2019 ; Handsaker *et al*., 2011). In short, this cannot distinguish between intact and truncated copies between accessions at a given locus. Quadrana *et al*. added to the split-reads approach a validation based on TSD (target site duplication) generated by most TE families upon insertion (Quadrana *et al*., 2016). But this method cannot be used in TEs which are devoid of TSD such as Helitrons. panREPET aims to analyse precisely all types of TEs. The TEMP pipeline (Zhuang *et al*., 2014 ; Stritt *et al*., 2018) combines PEM and split-read methods. By comparing panREPET against TEMP tool (Results - Benchmarking panREPET), in a subset of copies covering at least 80% of their consensus, we saw that panREPET detects globally many more shared TE insertions (TEMP retrieved only 396 copies among 24,085 detected by panREPET (1.6%), **Table 2**). Furthermore, TEMP tends to overestimate the number of accessions sharing a TE insertion while panREPET underestimates (**Table 2**). Our bidirectional detection method may be more stringent allowing eliminations of false positives.

Short reads identify position but do not give access to the exact sequence composition. A TE copy is generally about 300 bp to 10 kbp long, which is often larger than the average short-read length of about 100-300 bp (Groza *et al*., 2023). However, for TE regulation analysis, we need to know the exact activator and repressor sequence on the TE (Qiu *et al*., 2020). Some methods as x-Transposable element analyzer (xTea) overcome the short read limitation by including long reads (Chu *et al*., 2021). An additional problem is that read alignments for sequences of genomes to an assembled reference genome do not access the genetic environment of TE insertions in the genome from which the reads come. xTea tool tried to overcome this limit by using the Linked-Read technology from 10X Genomics. This utilizes microfluidics to partition and barcode DNA fragments from the same region, so that long-range information is embedded in the short-read data (Chu *et al*., 2021). Plus, thanks to long-read data, xTea utilizes the fully reconstructed TE sequences to provide additional information that cannot be obtained from short-read data (Chu *et al*., 2021). However, our approach is straightforward and does not involve any complex sequencing strategy as it works on several *de novo* assembled genomes. Indeed, panREPET provides direct access to the sequence of the whole TE copy, its exact insertion length, and its exact genetic environment. For instance, in our core TE insertion analyses, we were able to intersect TEs against genes and TFBS and calculate the exact coverages.

By comparing panREPET with Minigraph (Li *et al*., 2020) (see Results - Benchmarking panREPET), we observed that about 40-50% of TE copies detected by panREPET are retrieved by Minigraph. The remaining undetected TE copies highlight the limitation of detecting SVs before annotating the TE copies. We see that this kind of approach can lead to the detection of partial TE copies and when the TE copy is incompletely detected, Minigraph loses specificity. Indeed, a graph approach is more able to detect an insertion or deletion within the TE copy as a variant, but not the entire copy **(SF 6)**. Annotating TEs first is the guarantee of detecting the complete sequence. GraffiTE (Groza *et al*., 2024) uses several assembled genomes from the same species. It aligns each assembled genome against the reference but without aligning the others pairwise. It first detects structural variants (SVs) using genome-graphs and then annotates them in TEs. We have chosen a different paradigm: annotate the TEs in each genome independently and then compare TE copies pairwise between accessions. By comparing panREPET with GraffiTE (see Results - Benchmarking panREPET), we observe that GraffiTE detects better unique and cloud TE copies than shell and soft-core TE insertions **(Supplementary Table 4)**. GraffiTE tends to overestimate the number of individuals sharing a TE insertion **(Supplementary Table 4)**. Finally, Minigraph and GraffiTE are reference centric method to detect structural variants, while panREPET is designed to detect variations of TE insertions without depending on a reference. Indeed, panREPET extracts copies for each assembly that may have undergone an insertion or deletion relative to their consensus at the TE copy scale. panREPET provides a description of the nucleotid preservation on the TE sequence after its insertion.

When assemblies are at a chromosomal scale, we are able to distinguish a translocation event or a case of segmental duplication (paralog copies, data not shown). We can identify a translocation occurring after the TE copy insertion whether a shared TE copy is present in a chromosome but an other TE copy from an other individual is present in an other chromosome. Thanks to our bidirectional detection approach, we can also highlight a segmental duplication if for a shared TE copy, an other individual sharing the copy presents two best hits : the first one in the same chromosome and the other one in other chromosome.

### New biological results

A previous consensus library for *B. distachyon* was available from the TREP database and was composed of 233 sequences (Schlagenhauf *et al*., 2016, Stritt *et al*., 2019). This library contained 74% TIR, 20% LTR (10% Copia, 10% Gypsy) and 6% Helitron. Our new consensus library is more complete with 1,995 sequences including new TE families (LINE, SINE, MITE and TRIM) (**Fig. 2d**). The TE library was built *denovo* from the Bd21 reference (see Materials and methods - Genome analysis tools). MITEs and SINEs are the main component of core and soft-core pangenomic compartments (**Fig. 2d**), hence our TE library allows us to study ancient and conserved TE insertions that has not been possible before in other studies. MITEs with TFBS are conserved at the species level. Concerning core TE insertions conserved in the species but devoid of genes or TFBS, we observed that they seem to be conserved by the effect of a nearby gene under positive selection.

In *B. distachyon*, panREPET detected a majority of unique TE insertions (**Fig. 2a-b**). That is expected as we extracted full TE copies (or almost, see Results - panTEannot) and the number of unique TE insertions may depend on assembly quality **(SF 2)**. Interestingly, Stritt *et al*. suggested that *B. distachyon* expands populations without inter-accession breeding (Gordon *et al*., 2017, Stritt *et al*., 2020); this phenomenon may lead to an excess of singletons (Bourgeois *et al*., 2019).

Our approach aims to understand how individual TE lineages have evolved at the species level. A TE insertion shared by a subset of individuals could originate by two scenarios. First, the TE insertion could be inherited only by closely related accessions as it is recent. Second, the TE insertion could be lost by some accessions over time through excision, ectopic recombination, genetic drift or purifying selection. To distinguish these possibilities, we can estimate TE insertion age. Usually, a molecular clock of 1.3 × 10^-8^ substitutions/site/year is used on LTR-retrotransposons (SanMiguel e*t al.*, 1998). However, it does not take into account the evolutionary history of the species and it does not consider all types of TEs. Stritt *et al*. dated TE insertions among these 54 genomes of *B. distachyon* but only for LTR copies (Gordon *et al*., 2019). On the contrary, panREPET aims to date TE insertions for all types of TE copies thanks to the identification of shared insertions between individuals. In our study, a genetic tree based on whole-genome SNPs was available (Gordon *et al*., 2017) as well as genetic cluster dating based on a coalescence method (Minadakis *et al*., 2023) (see Results - TE insertion Dates). Hence, by fetching pairwise genetic distances among accessions, we were able to date TE insertions by considering genome-wide evolution and the specific events undergone by the species such as bottlenecks.

We dated the TE insertions before and after the Ice Age. We detected four ancient bursts of transpositions in *B. distachyon*: (i) at least 45 kya before or during the split into three lineages. However, it should be noted that panREPET utilizes assembled genomes as an input, a requirement that does not align with the characteristics of Italian genomes. Their addition would enable even more precise dating, as it would be the ancestral clade (Stritt *et al.,* 2022; Minadakis *et al*., 2023); (ii) 35 kya which could correspond to the bottleneck undergone by the S+ genetic cluster; (iii) 22 kya during the Last Glacial Maximum (LGM) and occurring after the abrupt population effective size decline around 30 kya; (iv) and during the Holocene around 10 kya with the recolonization of Europe and the Middle East.

After the Holocene, the accessions Bd21 and Bd21-3 from arid climate seem to have activated specific TEs potentially induced by abiotic stresses like pollution or drought (**Fig. 6c’)**. Due to the restriction of TFBS data availability to Bd21, there is a paucity of information regarding copies that are not shared with Bd21. Moreover, a homogeneous distribution of accessions by climate class would enable more precise inference of local activations, although in the end functional validations of TEs likely to be involved in adaptation will be required. panREPET aims to facilitate this type of automated query for a large dataset.

## Conclusion

We developed and have presented here a new bioinformatic tool called panREPET that makes it possible to retrace TE insertions in a species in a more accurate way than current approaches because: (i) it provides exact genomic coordinates of shared TE insertions for all individuals and their exact sequences without depending on a reference genome, which can provide a description of the nucleotid preservation after its insertion (ii) the comparison is made at the TE scale and not at the SV scale which permits to work at the TE copy scale (iii) it facilitates dating all types of TE copies where methods do not take into account the population dynamics (eg. population bottlenecks or expansion), and the most used methods just allow dating for LTR families). Finally, we dated two major TE bursts corresponding to major key climate events and found specific TE families that seem to be activated in arid climate induced by pollution or drought after the Holocene.

## Materials and methods Brachypodium distachyon data

We downloaded fifty-three whole-genome *de novo* assemblies *of B. distachyon*, their gene annotations and their gene ontology come from the Phytozome database (https://phytozome.jgi.doe.gov/) (Gordon *et al*., 2017). The BdTR7a genome comes from Stritt *et al*., 2019. The genome versions are detailed in **Supplementary Table 1**.

We downloaded 17,589 predicted TFBS in promoter regions (−500 bp ∼ +100 bp of transcription start site) and 144,761 predicted TFBS genome wide from the PlantRegMap database which uses the FunTFBS algorithm (https://plantregmap.gao-lab.org/download.php) (Tian *et al*., 2019). Data are only available for the reference Bd21. We inferred the gene ontology associated to this TFBS thanks to a summary of annotation details available from Phytozome (Bdistachyon_556_v3.2.annotation_info.txt).

We fetched centromeric coordinates on Bd21 v3 from Li *et al*., 2018, on Table 1 (Li *et al*., 2018). The recombination rates are from Huo *et al*., 2011, after mapping data to the version 3 reference genome assembly. We linked to the version 3 by searching for the sequences flanking the polymorphic SNPs in the genome version 3 to get the coordinates.

We extracted whole-genome SNP distances from phylogenetic tree in Newick format from Gordon et *al.*, 2017, Supplementary Figure 4.a. We converted newick format to pairwise distances between accessions, using the Phylo module from Biopython v3.10.13 (Cock *et al*., 2009).

We used the kgc R package v1.0.0.2 (Beck *et al*., 2018) to associate each *B. distachyon* accession location with a general climate by converting the geographic coordinates into decimal degrees.

### Genome analysis tools

We built a *de novo* TE library from the reference Bd21 v3.2 with the TEdenovo pipeline (Flutre *et al*., 2011) from REPET v3.0 (https://urgi.versailles.inrae.fr/Tools/REPET). The clustering step was done with GROUPER (Quesneville *et al*., 2003) only. The TE library is composed of consensus sequences which are reference sequences. We curated the annotation automatically by a second TEannot process (Jamilloux *et al*., 2017) reducing the number of consensus sequences from 4475 to 1,995. The TE classification comes from the classification of their consensus performed by the PASTEC classifier from REPET package v3.0 (Hoede *et al*., 2014). Some consensus sequences were classified by inferring their classification with consensus sequences belonging to the same cluster (or MCL).

We calculated sensitivity and specificity as in Baud *et al*. (Baud *et al*., 2022) ; True positives (TP) are defined as predicted TE nucleotides that truly belong to a TE copy. False positives (FP) are the predicted TE nucleotides that do not really belong to a TE copy. True negatives (TN) are the nucleotides correctly predicted not to belong to a TE copy (correct rejection), and false negatives (FN) are the true TE copy nucleotides missed by the TE prediction process. Sensitivity, the true positive rate, given by the formula TP/(TP+FP), is obtained by calculating the fraction of nucleotides in the predicted TE overlapping with the TE reference annotation. Specificity, also refered to as the true negative rate, is less straightforward to calculate. It can be calculated according to the formula TN/(TN+FP), but TN and FP are difficult to determine for TEs, as they can only be known if we are sure that we have identified all the TE copies in the genome, which does not really seem possible. However, as a first approximation, we can consider that genes are not TEs, and are not derived from TEs, and use this information to obtain more accurate estimates for TN and FP. This is obviously an approximation as TEs are known to be sometimes part of genes. Hopefully this is rare compared to other regions of the genome, in particular if we consider the coding sequence (CDS) as it discards introns as well as 5’ and 3’ UTR where TEs can be found frequent. In this context, FP are predicted TE nucleotides that overlap a gene CDS annotation, and TN are CDS regions not predicted to be TEs. Accuracy, given by ACC=(TP+TN)/(TP+TN+FN+FP), is the rate of correct predictions.

We annotated the fifty-four genomes of *B. distachyon* with the TEannot pipeline from REPET v3.0 without mreps (Quesneville *et al*., 2005).

The Bedtools version is 2.30.0. *bedtools intersect* is configured with option –wo writing the original A and B entries plus the number of base pairs of overlap between the two features (Bioinformatics, 2024).

We built clustermaps with Seaborn package from Python v3.10.13 (Waskom, 2021).

We performed ANOVA with statsmodels module from Python v3.10.13 (Cock *et al*., 2009). We performed T test with SciPy library from Python v3.10.13 (Virtanen *et al*., 2020).

### Benchmarking panREPET

In order to facilitate a comparison with the TEMP results of Stritt *et al*. (2018), we conducted a panREPET analysis on the same set of 53 genomes. For this analysis, we utilized version 2.0 of the *B. distachyon* reference genome. The *B. distachyon* TE consensus sequences were fetched from the TREP database (http://botserv2.uzh.ch/kelldata/trep-db/index.html), as reported by Stritt *et al*. The coverage between TE copies and their consensus was configured at 80%. For comparing with Minigraph and GraffiTE, we used panREPET results as Results - Retracing shared TE insertions i.e. on the 42 *B. distachyon*.

Minigraph heavily depends on the first genomes introduced as its graph construction consists in adding genomes incrementally. Hence, we sorted the input genomes in descending order of assembly quality (we used the assembled genome size after removing ‘N’ occurrences which is a a compromise between N50 contigs and N50 scaffolds, see Results - panTEannot). We submitted as the first genome BdTR7a in PacBio because it is the only genome in long read among dataset. Minigraph was configured by default including only SVs affecting 50 bp of sequence or more.

GraffiTE was used with the default parameters.

We used *bedtools intersect* from Bedtools v2.30.0 for intersecting coordinates TE insertions between TEMP or Minigraph or GraffiTE results against panREPET results. To calculate the proportion of individuals in common between two shared TE insertions detected by two different tools, we fetched the number of same individuals which is then divided by the largest number of individuals sharing the TE insertion over these two TE insertions.

## Data availability

The *Brachypodium distachyon* (Bd21) TE library and TE annotation file (gff) are available in the RepetDB (Amselem *et al*. 2019) database (https://urgi.versailles.inrae.fr/repetdb/begin.do).

## Code availability

panTEannot source code is available at https://forgemia.inra.fr/urgi-anagen/panTEannot. panrREPET source code is available at https://forgemia.inra.fr/urgi-anagen/panREPET.

## Acknowledgement

We thank Maela Semery for providing climate classes of each accession (see Materials and Methods - Brachypodium distachyon data). We thank Clémentine Vitte, Leandro Quadrana and Anne Roulin for their relevant comments. We thank Gautier Sarah for testing panTEannot.

## Author information - Contributions

HQ and JC conceived the project. MB, MRO and SS developed panREPET. SS developed panTEannot and generated all manuscript results on *B. distachyon*. SS wrote the manuscript under the supervision of HQ and JC.

