## Supplementary Information for "panREPET: a reference-free pipeline for detecting shared Transposable Elements from pan-genomes to retrace their dynamics in a species"

### Supplemental Figures

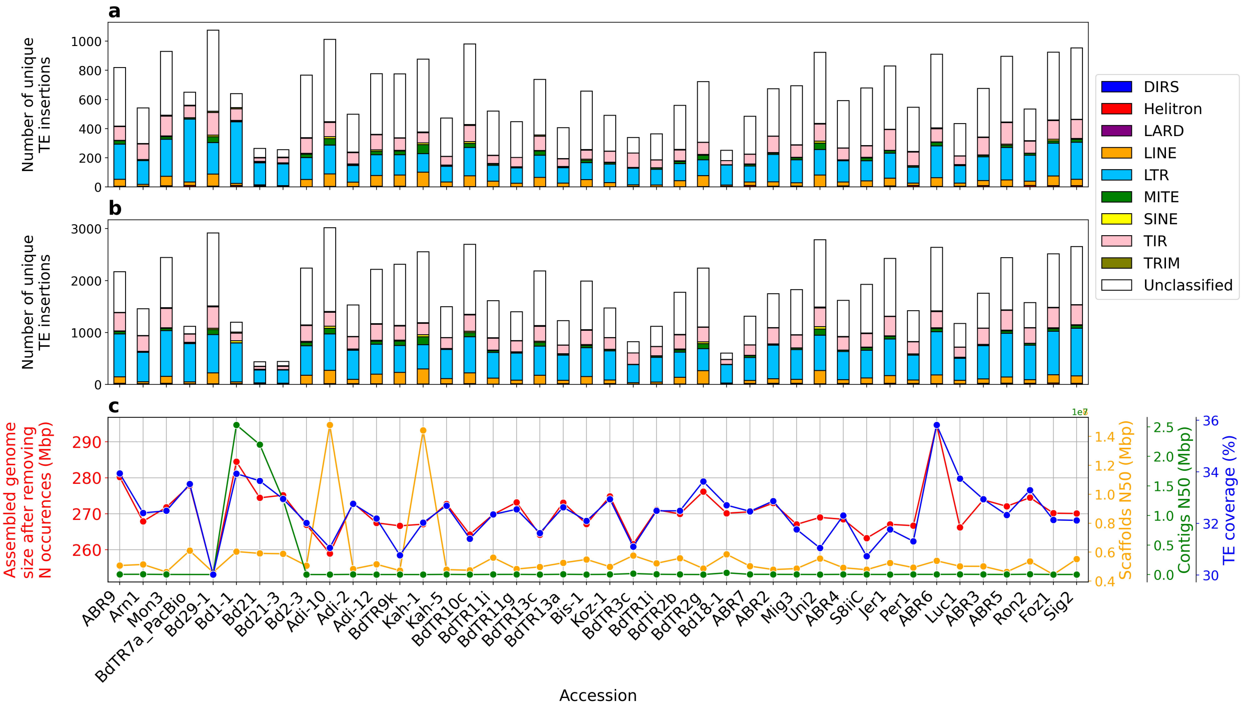


**Supplementary Figure 1: a, b** Distribution of the number of unique TE insertions among accessions. Accessions are ordered as the whole-genome SNP genetic tree (Gordon *et al.*, 2017, Supplementary Figure 4.a). **a** The percentage coverage between the TE copy and its consensus (parameter covcons) is 95-105%. **b** covcons=75-125% **c** Assembled genome size after removing N occurences (Mbp), Scaffolds N50 (Mbp), Contigs N50 (Mbp) and TE coverage from panTEannot (%) of each accession.

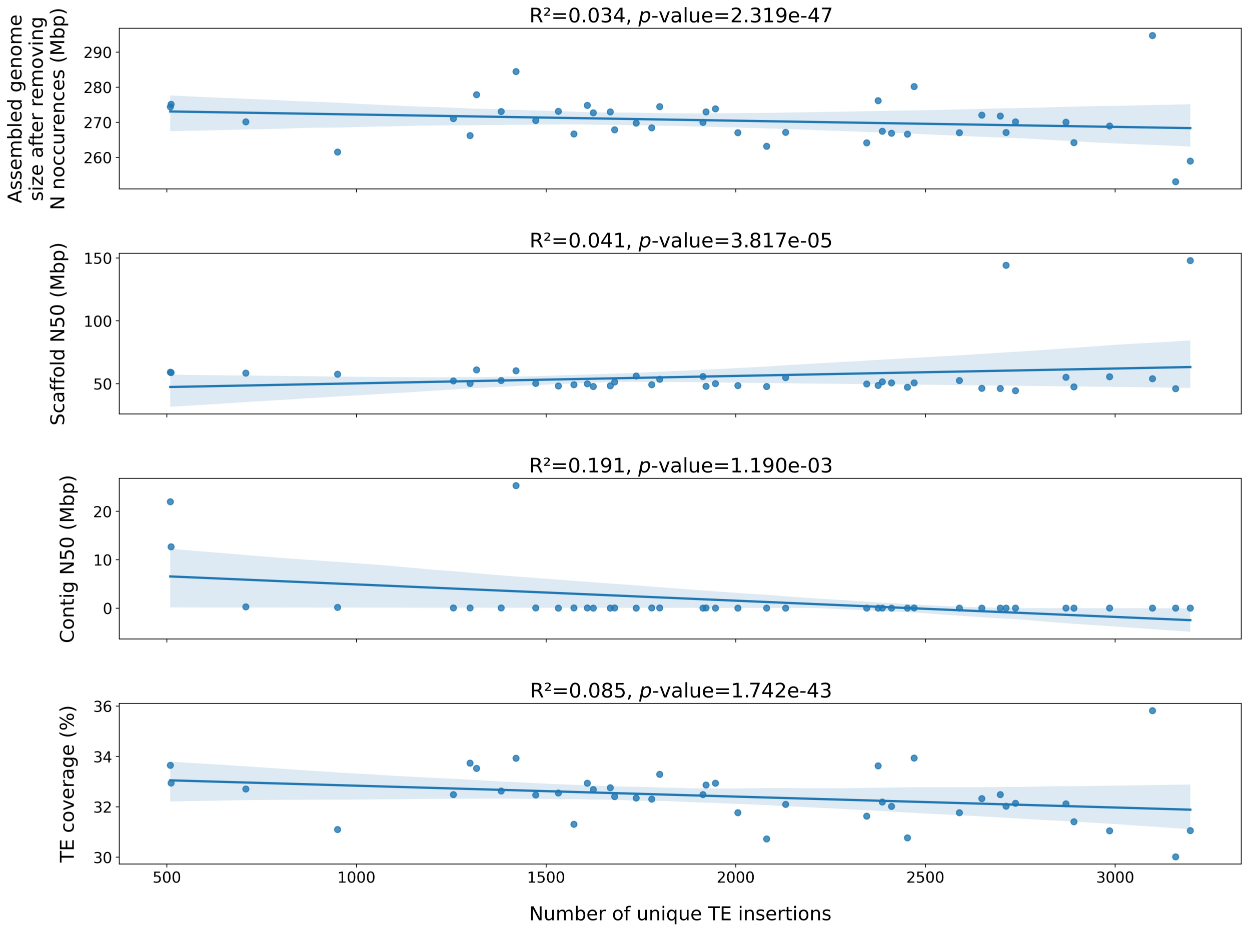


**Supplementary Figure 2:** Four scatterplots comparing linear regressions for dependent variable number of unique TE insertions with four explanatory variables (in order): Assembled genome size after removing N occurences $R^2$=0.034, *p*-value=2.319e-47 ; Scaffolds N50 $R^2$=0.041, *p*-value=3.817e-05 ; Contigs N50 $R^2$=0.191, *p*-value=1.190e-03 and TE coverage $R^2$=0.085, *p*-value=1.742e-43. The percent coverage between copy and its consensus parameter here is 75-125%.

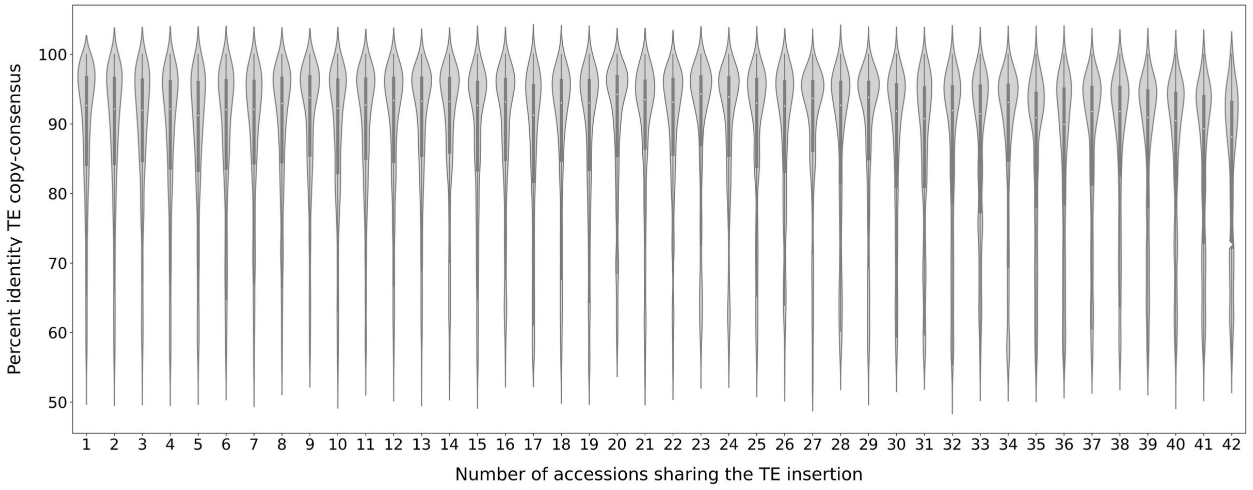


**Supplementary Figure 3:** Boxplots showing the age distribution of TE insertions according to the number of accessions sharing it. The age is estimated by percentage identity between the TE copy and its consensus.

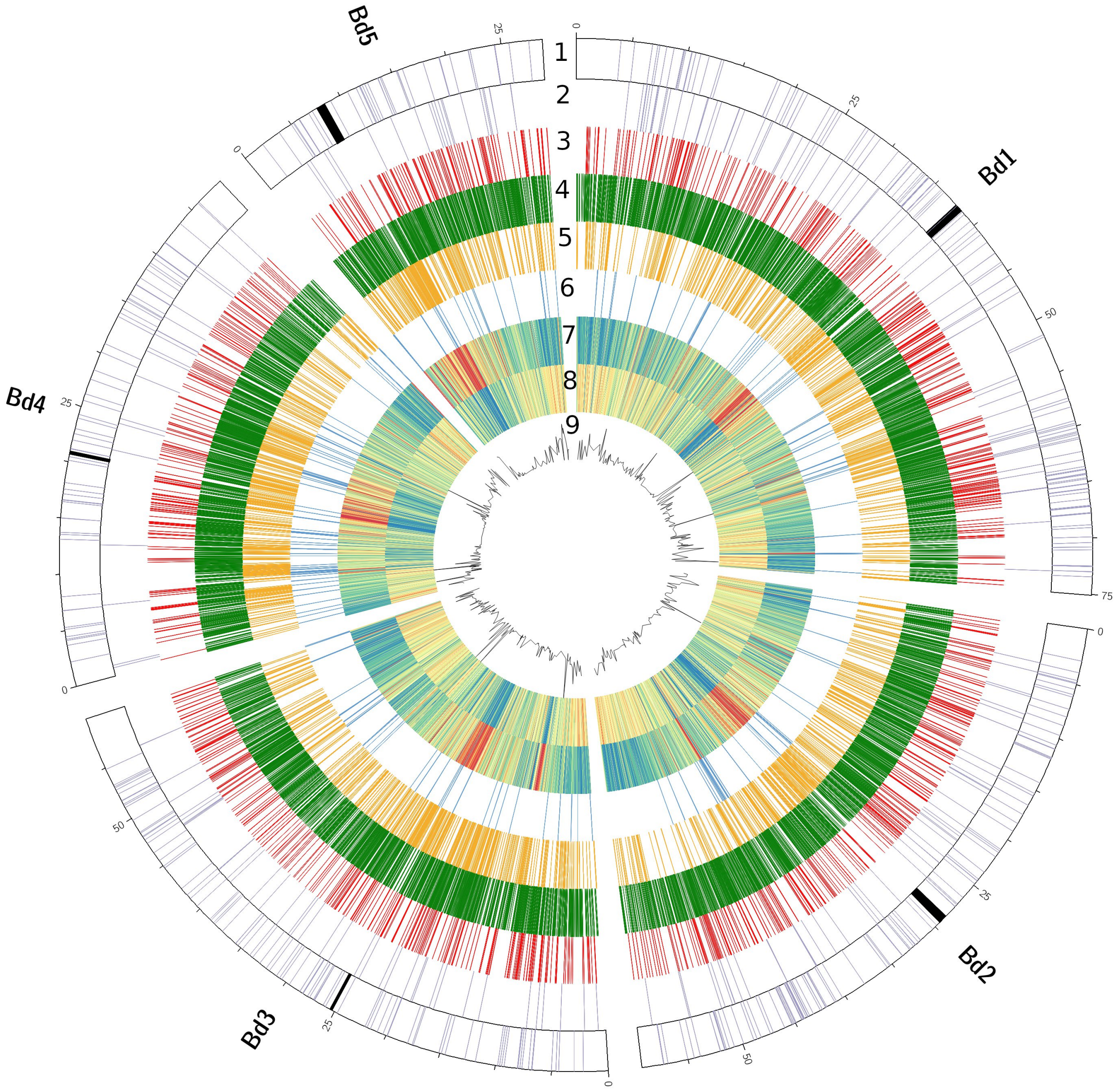


**Supplementary Figure 4: (Layers 1-6)** Distribution among chromosomes of TE insertions. **(Layer 1, outer layer)** *Not gene related core TE* insertions. Centromeres are in black. **(Layer 2)** Core TE insertions intersecting a gene or TFBS. **(Layer 3)** Soft core. **(Layer 4)** Shell. **(Layer 5)** Cloud. **(Layer 6)** Unique. **(Layer 7)** The TE density is from blue to red (range 1-99%) and was calculated per 50 kbp interval. **(Layer 8)** The gene density is is from blue to red (range 0-84%) and was calculated per 50 kbp interval. **(Inner layer)** Recombination rates. Chromosome visualization was plotted by circos-0.69-9.

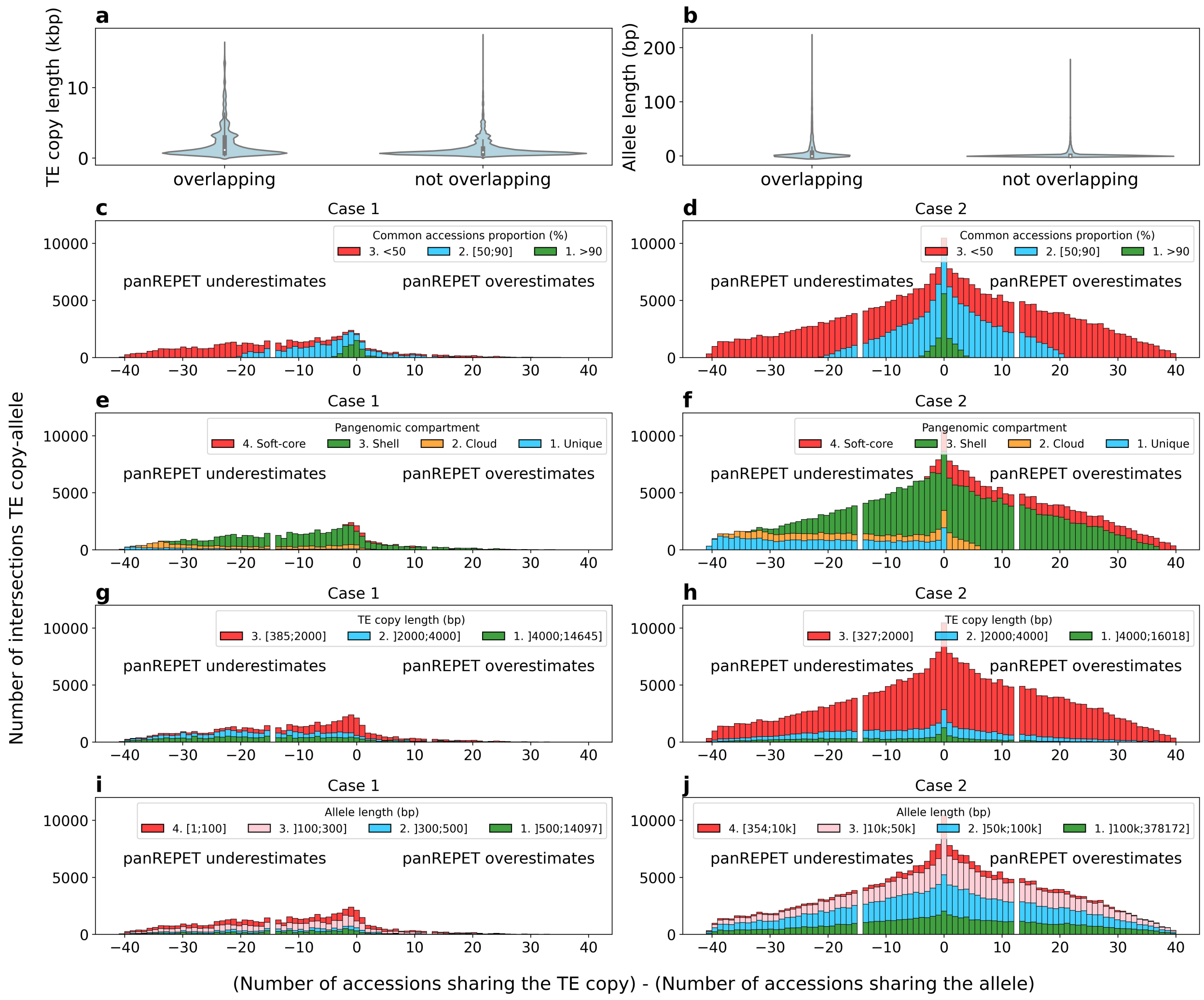


**Supplementary Figure 5: (a)** TE copies lengths from panREPET (kbp). **(b)** Allele lengths from Minigraph (bp). **(c-j)** Histogram of differences between the number of accessions sharing the TE copy from panREPET and those sharing the associated allele from Minigraph. **(c,e,g,i)** The allele is included in the TE copy (case 1). **(d,f,h,j)** The TE copy is included in allele (case 2). **(c,d)** The colors represent the proportions of accessions that are the same. **(e,f)** The colors represent the pangenomic compartment. **(g,h)** The colors represent the TE copy lengths **(i,j)** The colors represent the allele lengths. For readability, we have not shown the results for covcons=75-125% (percentage of coverage on the consensus of the TE copies), but the observations are the same.

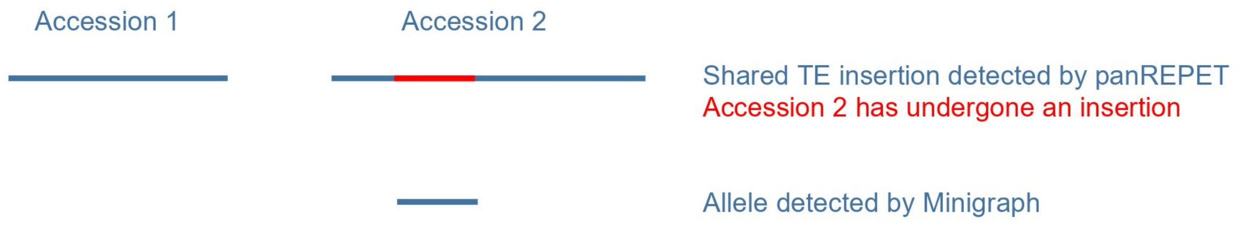


**Supplementary Figure 6:** Alleles from Minigraph may correspond to fragmented TE copies

#### Supplemental Tables

**Supplementary Table 1 :** Accession information (Genome source: https://phytozome-next.jgi.doe.gov/info/[Phytozome_genome_ID]) and their TE coverages. Genomes removed from the study are in bold and marked with an asterisk. The reference genome is in bold.

**Supplementary Table 2 :** Sheet 1: List of traits describing the 42 accessions from *B. distachyon*. Color codes correspond to the colors used in **Figure 6**. Sheet 2: Climate traits for each accession, in raw format.

| | TEMP | | panREPET |
|---|---|---|---|
| | **TAPs** (Bd21 is present) | **TIPs** (Bd21 is absent) | **Cliques** |
| Number of shared TE insertions | 1,889 | 3,627 | 70,373 TE copies in Bd21 genome annotated by panTEannot<br>24,492/70,373 TE copies in Bd21 covers their reference on more than 80% (34%)<br>44,597 cliques including 24,085 TE copies shared with the reference Bd21 (equivalent to TAPs) and 646 unique Bd21 TE copies |
| TE annotations retrieved by panTEannot | 1,204/1,889 (63.7%) | NA | / |
| Select TAPs that intersect TE copies in Bd21 annotated by panTEannot and covering their reference on more than 80% | 396/1,204 (32%) | NA | / |
| Number of retrieved TE insertions | 396/1,204 (32%) | 118/3,627 (3.2%) | 396/24,085 (1.6%) |
| Number of accessions (TEMP) minus Number of accessions (panREPET) per common TE insertion | Mean: 25<br>Median: 25 | Mean: -34<br>Median: -42 | Mean: 25<br>Median: 25 |
| Proportion of identical accessions | Mean: 39.4%<br>Median: 38.2% | Mean: 5.87%<br>Median: 2.19% | Mean: 39.4%<br>Median: 38.2% |
| **TE lengths (bp)** | | | |
| Common TE library from TREP database (233 sequences) | Min: 84 ; Mean: 4,525 ; Median: 3,951 ; Max: 15,485 | | |
| All annotated TE copies | Mean: 951 (Stritt *et al.*, 2018) | NA | (Bd21 TE annotation)<br>Mean: 942<br>Median: 175 |

**Supplementary Table 3 :** Comparison between TEMP and panREPET tool

<table>
<tr><th colspan="2"></th><th colspan="2">GraffiTE</th><th>panREPET</th></tr>
<tr><th colspan="2"></th><th>Deletion<br>(Bd21 is present)</th><th>Insertion<br>(Bd21 is absent)</th><th>Cliques + unique TE copies</th></tr>
<tr><td colspan="2">SVs containing TEs</td><td colspan="2">47,236</td><td>/</td></tr>
<tr><td colspan="2">Total number of TE copies</td><td>111,860</td><td>48,241</td><td>/</td></tr>
<tr><td colspan="2">Number of TE copies/SV</td><td>Mean: 3.8<br>Median: 1</td><td>Mean: 2.6<br>Median: 2</td><td>/</td></tr>
<tr><td colspan="2">Number of accessions per SV (GraffiTE) or per TE insertion (panREPET)</td><td>Mean: 1<br>Median: 1<br>Max: 3</td><td>Mean: 1<br>Median: 1<br>Max: 11</td><td>Mean: 19<br>Median: 18<br>Max: 41</td></tr>
<tr><td colspan="2">SVs containing TEs covering their consensus on more than 95% and less than 105%</td><td colspan="2">4,314/47,236 (9.1%)</td><td>/</td></tr>
<tr><td colspan="2">Total number of TE copies covering their consensus on more than 95% and less than 105%</td><td>6,487/111,861 (5.7%) corresponding to 3,166 shared TE insertions with Bd21</td><td>1,169/48,241 (2.4%)</td><td>17,918 cliques whose 9,192 TE copies shared by the reference Bd21<br>+<br>282 unique TE copies</td></tr>
<tr><td colspan="2">Number of retrieved shared TE insertions</td><td>1,802/3,166 (56%)</td><td>23/1,169 (1.9%)</td><td>1,802/9,474 (19%)<br>Details :<br>Soft-core : 65/1,098 (5.9%)<br>Shell : 965/5,966 (16%)<br>Cloud : 679/2,240 (30%)<br>Unique 93/282 (32%)</td></tr>
<tr><td colspan="2">Number of accessions (GraffiTE) - Number of accessions (panREPET) per shared TE insertions</td><td>Min: -5<br>Mean: 26<br>Median: 30<br>Max: 40</td><td>/</td><td>Min: -5<br>Mean: 26<br>Median: 30<br>Max: 40</td></tr>
<tr><td colspan="2">Proportions of identical accessions (%)</td><td>Mean: 33<br>Median: 24</td><td>/</td><td>Mean: 33<br>Median: 24</td></tr>
<tr><td colspan="5">Sequence lengths (bp)</td></tr>
<tr><td colspan="2">Common TE library (1,195 sequences)</td><td colspan="3">Min: 335<br>Mean: 2,410<br>Median: 1,285<br>Max: 17,319</td></tr>
<tr><td rowspan="3">TE copies covering their consensus on more than 95% and less than 105%</td><td>All</td><td>Mean: 3,234<br>Median: 2,614</td><td>Mean: 3,567<br>Median: 3,052</td><td>Shared : Mean: 2,323 ; Median: 1,258<br>Unique: Mean: 6,369 ; Median: 5,452</td></tr>
<tr><td>Retrieved</td><td>Mean: 3,383<br>Median: 3,005</td><td>Mean: 3,326<br>Median: 3,113</td><td>Mean: 3,184<br>Median: 2,697</td></tr>
<tr><td>Not retrieved</td><td>Mean: 3,271<br>Median: 2,476</td><td>Mean: 3,326<br>Median: 2,611</td><td>Mean: 2,098<br>Median: 1,089</td></tr>
</table>

**Supplementary Table 4 :** Comparison between GraffiTE and panREPET tools

### Supplementary Results

**Detailed panTEannot pipeline**

We used Blaster (version 2.31) from the Singularity container te_finder_2.31.sif (https://cloud.sylabs.io/library/hquesneville/default/te_finder) to align chunked genomes against the consensus sequences from TE library. The cutterDB step was configured as -l 200000 -o 10000, meaning that chunks have a length of 200kbp and they overlap over 10kbp.

Matcher (version matcherThreads2.31) is called up from the Singularity container te_finder_2.31.sif (https://cloud.sylabs.io/library/hquesneville/default/te_finder). We used the option join (-j) to join fragments that it considers close enough to represent an insertion or deletion. We use the cleanup after join option (-x) to clean up fragments overlaps.

**Detailed panREPET pipeline**

*About Bidirectional best-hit detection (Step 3):*

We used Minimap2 version 2.24-r1122. The long assembly to reference mapping is up to 5% sequence divergence (-asm20) to be sensitive. The score alignment provided by Matcher is used to select the best hit between a pairwise comparison of all copies of the accessions.

We save computing time by only comparing TE copies annotated from the same consensus, and it is possible to compare only TE copies present on the same chromosome when the assembly is at the chromosome level.

A trade off between the flanking region length and the identity is expected on the flanking region. Preliminary analysis on 8 *Arabidopsis thaliana* genomes (Col0, Ler0, KBSMac74, Nd1, Ler1, Bur0, C24, Kro0) showed that the larger the extension size is, the greater the number of detected shared TE insertions are. We observed that using a more significant extension length (2000 bp) can cause a loss in sensitivity because larger flanking regions can contain repetitive sequences or other TE insertions. For this reason, we run the pipeline with an extension of the flanking region of 500 bp.

We also applied two filters to compare their effectiveness in the discovery of false positives: (i) an initial filter over the coverage percentage of the copy plus its flanking regions regarding its match (sequence over which the copy was aligned). However, our preliminary analysis on the 8 *Arabidopsis thaliana* genomes showed that this filter might produce false positives with some match occurring only in the flanking extremities and not in the copy’s sequence. (ii) We tested an another filter controlling a minimum threshold of coverage of the copy and their flanking extremities. This filter diminishes the number of false positive copies found only with the first filter. Finally, we only used a flanking coverage filter (80% by default) as partial copies with internal insertion/deletion can be hence selected.
